# A tripartite bacterial-fungal-plant symbiosis in the mycorrhiza-shaped microbiome drives plant growth and mycorrhization

**DOI:** 10.1101/2023.07.19.549792

**Authors:** Changfeng Zhang, Marcel G. A. van der Heijden, Bethany K. Dodds, Thi Bich Nguyen, Jelle Spooren, Alain Held, Marco Cosme, Roeland L. Berendsen

## Abstract

Plant microbiomes play crucial roles in nutrient cycling and plant growth, and are shaped by a complex interplay between plants, microbes, and the environment. The role of bacteria as mediators of the 400-million-year-old partnership between the majority of land plants and, arbuscular mycorrhizal (AM) fungi is still poorly understood. Here we test whether AM hyphae-associated bacteria influence the success of the AM symbiosis. Using partitioned microcosms containing field soil, we discovered that AM hyphae and roots selectively assemble their own microbiome from the surrounding soil. In two independent experiments, we identified several bacterial genera, including *Devosia*, that are consistently enriched on AM hyphae. Subsequently, we isolated 144 pure bacterial isolates from a mycorrhiza-rich sample of extraradical hyphae and isolated *Devosia* sp. ZB163 as root and hyphal colonizer. We show that this AM-associated bacterium synergistically acts with mycorrhiza on the plant root to strongly promote plant growth, nitrogen uptake, and mycorrhization. Our results highlight that AM fungi do not function in isolation and that the plant-mycorrhiza symbiont can recruit beneficial bacteria that support the symbiosis.

## Background

The evolution of the mycorrhizal symbiosis is thought to have been an essential step that enabled the development of land plants 500 million years ago [1]. Arbuscular mycorrhizal (AM) fungi live in symbiosis with 80% of terrestrial plants [2] and help plants to access distant water and nutrient sources [3-9], facilitating plant adaptation to environmental change [10]. AM extraradical hyphae extend from plant roots and enlarge the host plant’s area of nutrient uptake. Plants, however, simultaneously interact with many microbes in addition to AM fungi, especially on the roots where the plant microbiome is dense and diverse [11, 12].

Also non-mycorrhizal members of the plant microbiome can strongly affect plant growth [11]. Some detrimental microbes invade the plant and cause disease. Others promote plant growth, either directly e.g., by providing nutrients, or indirectly by protecting the plants from pathogens and other detrimental microbes [13]. Plants, therefore, foster and shape a microbiome to their benefit by exuding a mixture of microbe stimulatory and inhibitory compounds [14, 15]. As a result, the rhizosphere, the zone of soil surrounding roots that is influenced by these exudates, typically constitutes a dense microbial community that is distinct from that of the surrounding bulk soil and is selectively assembled by the plant [11].

Similar to plants, AM fungi have been shown to interact with their surrounding microbes [16]. For instance, the soluble exudates of the AM fungus *Rhizophagus irregularis* can have either antagonistic or stimulatory effects on individual fungal and bacterial isolates [17]. Interestingly, there is even a symbiotic footprint of the plant microbiome as plants hosting AM fungi harbour a different microbiome compared to non-mycorrhizal plants [18]. It has therefore been argued that AM hyphae extend the rhizosphere with a hyphosphere in which they similarly selectively assemble a microbiome [19].

Interactions between AM fungi and the microbes have primarily been studied by *in vitro* experiments, and have, e.g., revealed that bacteria can have different affinity for mycorrhizal hyphae [20, 21]. In recent years, some *in situ* experiments have been also conducted where soil with AM hyphae was compared to soil from which AM fungi were restricted. Through amplicon sequencing, these studies have shown that the bacterial community in soil with AM hyphae differed significantly from that of the bulk soil [22, 23]. A high throughput stable isotope probing research found that specific bacterial phyla attached to AM hyphae assimilated the most AM fungi-derived carbon [24]. Moreover, a recent study revealed that mycorrhiza-mediated recruitment of complete denitrifying *Pseudomonas* bacteria reduces N_2_O emissions from soil [25]. These findings suggest that the interactions between bacteria and AM fungi play a crucial role in shaping the hyphosphere microbiome.

The interactions between AM fungi and bacteria do not only have an impact on the bacterial community but also greatly influence the performance of the AM fungi. The functioning of the mycorrhizal symbiosis depends on microbial communities in soil and some soils have been characterized as mycorrhiza suppressive soils due to inhibitory effects of specific microbes [26]. Nonetheless, mycorrhiza helper bacteria of diverse taxonomy were found to promote germination of AM fungal spores, AM fungi establishment and subsequent colonization of plant roots [12, 27-29]. Moreover, phosphate-solubilizing bacteria have been shown to mineralize organic phosphorus (P) so that inorganic P can subsequently be absorbed by the AM mycelium [8, 30]. These findings suggest that specific components of the soil microbiome might benefit AM fungi and promote their growth and functioning.

Excessive fertilizer and pesticide use in conventional agriculture cause pollution and biodiversity loss [31, 32], while organic farming avoids these practices [33] and promotes soil biodiversity, with mycorrhizal fungal species identified as keystone taxa [34, 35]. Although organic farming typically results in lower crop yields than conventional practices, understanding the soil microbiome and key players like AM fungi and its associated microbiome can improve sustainable agricultural practices and close this yield gap.

We therefore investigated the role of AM fungi in shaping soil microbiomes. In a first set of experiments, we grew plants in compartmentalized microcosms using soil from a long-term field experiment with conventionally and organically managed agricultural plots. We sampled root, hyphae, and soil from distinct compartments of the microcosms, and isolated hyphae-adhering bacteria. Using ITS and 16S amplicon sequencing, we identified and isolated specific bacterial genera that are consistently enriched in hyphal samples. In a next set of experiments, we tested the effect of the AM fungi-associated bacterial isolates on plant performance. We discovered that *Devosia* sp., an AM fungi-associated bacterium, stimulated AM fungi colonization but also directly promoted plant growth by enhancing plant nitrogen (N) uptake.

## Results

### Experiment I: AM fungi-associated microbes on extraradical hyphae in a sterilized soil substrate

To understand the role of mycorrhizal hyphae in shaping the soil microbiome, we started by growing *Prunella vulgaris* (henceforth: Prunella) plants from a long term farming system and tillage (FAST) experiment at Reckenholz (Switzerland) that had either been managed with organic or conventional cultivation practices since the summer of 2009. Prunella is a common grassland plant in Switzerland, grows at the FAST trial location, and is regularly used as a model plant that strongly associates with, and responds to AM symbionts [31-36]. The plants were grown in the middle compartment of a 5-compartment microcosm (Fig. 1A). This middle compartment (COMP3) contained either organic or conventional soil (OS or CS) substrate, whereas the other compartments were filled with soil substrate to promote colonization of these compartments by extraradical AM hyphae. The compartments were separated by a 30-μm nylon filter that restrained the growth of roots inside the COMP3 but allowed extraradical hyphae to pass through and exit COMP3 into the compartments 4 and 5 (COMP4 and COMP5; Fig. 1A).

**Fig. 1.**
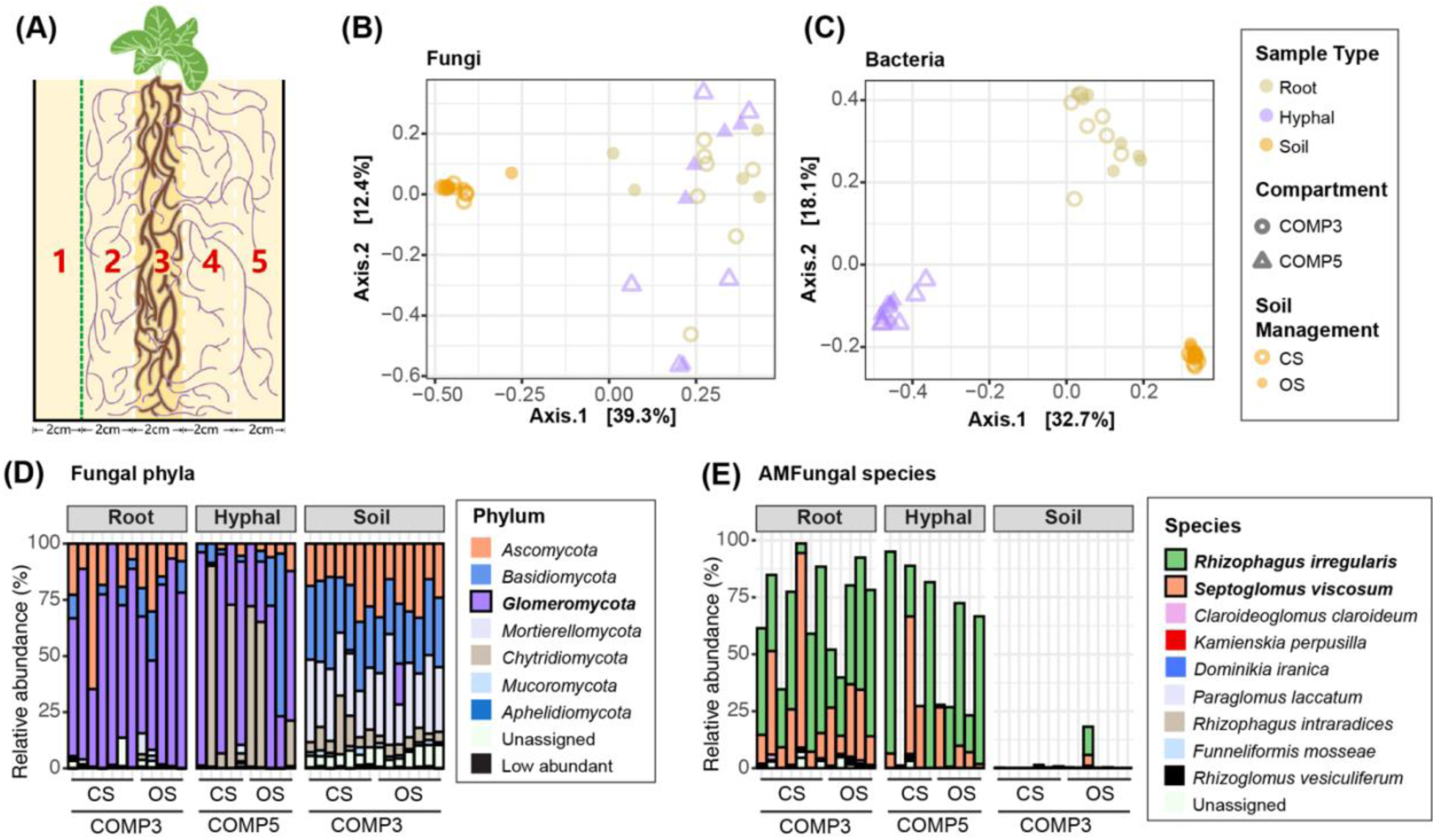
AM fungi-rich hyphal samples host a bacterial microbiome that is distinct from root and soil samples. **(A)** Schematic representation of 5-compartment microcosm in Experiment I. Compartment (COMP3) is filled with 30% of either organic (OS) or conventional (CS) soil, whereas COMP1, 2, 4 and 5 are filled with sterilized substrate. Roots are contained in COMP3 by filter mesh with 30-µm pores (white dashed lines), whereas extraradical AM hyphae are restricted from entering COMP1 by filter mesh with 1-µm pores (green dashed line). **(B)** PCoA of fungal communities using Bray-Curtis distances in root, soil and hyphal samples of plants growing in either CS (open symbols) or OS (closed symbols). **(C)** PCoA of bacterial communities in root, soil and hyphal samples of plants growing in either CS or OS. Colors in **(B)**, **(C)** indicate different sample types. Shapes depicts the compartments of microcosm. **(D)** Relative abundance of fungal phyla in root and soil samples from COMP3 and hyphal samples from COMP5. Colors represent the distinct phyla as indicated in the legend. Phyla with relative abundance below 1% were aggregated and categorized as low abundant. **(E)** Relative abundance of *Glomeromycota* spp. in root, soil and hyphal samples in Experiment I. Colors represent the distinct AM fungal species as indicated in the legend.

We cultivated the plants for 3 months, after which we found that extraradical hyphae had reached COMP5. We isolated DNA from these samples and subsequently analyzed the composition of fungal and bacterial communities by sequencing ITS and 16S amplicons, respectively.

#### Soil, roots, and hyphal samples represent distinct microbial communities

Principal coordinate analysis (PCoA) of the fungal communities showed a clear separation of soil samples from root samples and hyphal samples (Fig. 1B). Sample type explained a significant proportion (42.9%) of the variation within the fungal community, as determined by permutational multivariate analysis of variance (PERMANOVA; R^2^=0.429, F = 12.416, *p* < 0.001) and each of the sample types was significantly distinct from the two other sample types (Table S1). This shows that there is a significant rhizosphere effect shaping the fungal community on the root and that the hyphal samples consist of a fungal community that is slightly different from the root samples. In the 16S amplicon data, we observed a clear separation of bacterial communities between all sample types in the PCoA plot (Fig. 1C). Almost half (49.6%) of the variation is explained by sample type (PERMANOVA; R^2^=0.496, F = 18.751, *p* < 0.001) and a pairwise PERMANOVA test shows that all sample types (root, soil and hyphal) are significantly different from each other (Table S1). This shows that the hyphae picked from COMP5 harbor a bacterial community distinct from those in the root and soil samples. We hypothesized that the hyphal samples include the microbes that live around and attached to the mycorrhizal fungi, whereas the root samples additionally include those microbes that are promoted by the roots themselves.

#### Glomeromycota abundantly present in hyphal and root samples

*Glomeromycota*, the fungal phylum to which all AM fungi belong, were detected at 71% average relative abundance (RA) of the root fungal community, while on average 51% of the fungal reads in the hyphal samples of COMP 5 were annotated as *Glomeromycota*. *Glomeromycota* is thus the dominant fungal phylum in both the root and hyphal samples. In soil samples from COMP3, which were dominated by plant roots, however, this phylum was below 1% in 12 out of 14 samples (Fig. 1D). This shows that even in the FAST soil close to Prunella roots, AM fungi are lowly abundant, but that over the course of the experiment, AM fungi had colonized Prunella roots and had become very abundant on the roots. Moreover, AM hyphae had grown and extended from the roots in COMP3 to COMP5, where we were able to collect these hyphae using a modified wet sieving protocol. Within the *Glomeromycota*, we found sequences belonging to two prevalent AM species. *Rhizophagus irregularis* (average RA: 42% in root and 36% in hyphal samples, respectively) and *Septoglomus viscosum* (average RA: 25% in root and 14% in hyphal samples, respectively) were the most abundant species in the fungal community. In addition to *Glomeromycota, Chytridiomycota* also take up a considerable percentage of the reads in some of our hyphal and soil samples but were hardly detected on the roots. Hyphae of *Glomeromycota* cannot easily be distinguished from those of various other fungi, and consequently, a part of the collected hyphal samples belonged to non-mycorrhizal fungal species.

#### Effects of field management practices on soil microbiome negated on hyphae and roots

Previous work demonstrated that the soil microbiome is affected by soil management practices [35, 36]. The long-term FAST experiment contains plots that have been managed using either conventional or organic cultivation practices for over a decade. We filled microcosms with either FAST OS or CS soil to study the influence of management practices on the rhizosphere and hyphosphere microbiome composition. At the end of 3 months of Prunella cultivation in the greenhouse, the soil in COMP3 was still significantly influenced by preceding management practices of the FAST experiment. This is evidenced by a significant difference in the fungal and bacterial communities’ composition between OS and CS samples collected from the field (Fig. S1A, S2C; Table S2). We found that 4 fungal genera and 5 classes of bacteria were more abundant in OS, while 6 fungal genera and 2 bacterial classes were more abundant in CS (Fig. S1B, S1D; Table S2). Remarkably, we did not find significant effects of soil management on the microbiome composition in the root or hyphal samples of our Experiment I (Table S2). This suggests that the signature of soil management type on soil microbiome disappears while root and hyphae selectively assemble their microbiomes, even though the distinction of microbial communities between OS and CS can still be observed in the soil in between roots in COMP3 (Fig. S2). Moreover, the microbial difference between OS and CS soil affected neither mycorrhizal colonization nor plant performance (Fig. S3).

### Experiment II: Extraradical hyphae-associated microbes in non-sterilized soil substrate

In the experiment described above, we found that fungal hyphae from COMP5 harbor a microbial community that is distinct from the soil microbiome in COMP1 and the root microbiome in COMP3, the later containing the Prunella roots. However, these hyphae were collected from the sterilized soil substrate of COMP5 that was distinct from the soil substrate in COMP3. We followed up on this experiment by planting 2-week-old Prunella seedlings in the middle compartment (COMP3) of 5-compartment microcosms, but now we filled all compartments with the same non-sterilized OS substrate. Again, the roots were restrained to COMP3 by filters with 30-µm pore size that did allow extraradical growth of fungal hyphae to COMP4 and 5. Differently from Experiment I, we used in Experiment II filters with 1-µm pore size to prevent the growth of hyphae not only into COMP1 but also into COMP2 (Fig. 2A). We thus hoped to create compartments in each microcosm where the soil microbiome was shaped by the combination of root, hyphae, and their combined exudates (COMP3), by plant-associated hyphae alone (COMP5), or by neither roots nor hyphae (COMP1). We hypothesized that in addition to root COMP3, only buffer COMP2 and 4 would be affected by root exudates, of which COMP4 would additionally be shaped by the plant-associated hyphae that pass through them. After 3 months of Prunella cultivation, we sample soil from each of the compartments and in addition root samples from COMP3 and COMP5 hyphal samples. As we were unable to pick hyphae from unplanted microcosms, we were unable to obtain hyphal samples from unplanted microcosms, and we have to assume that most picked hyphae in the microcosms with Prunella plants belong to plant-associated fungi.

**Fig. 2.**
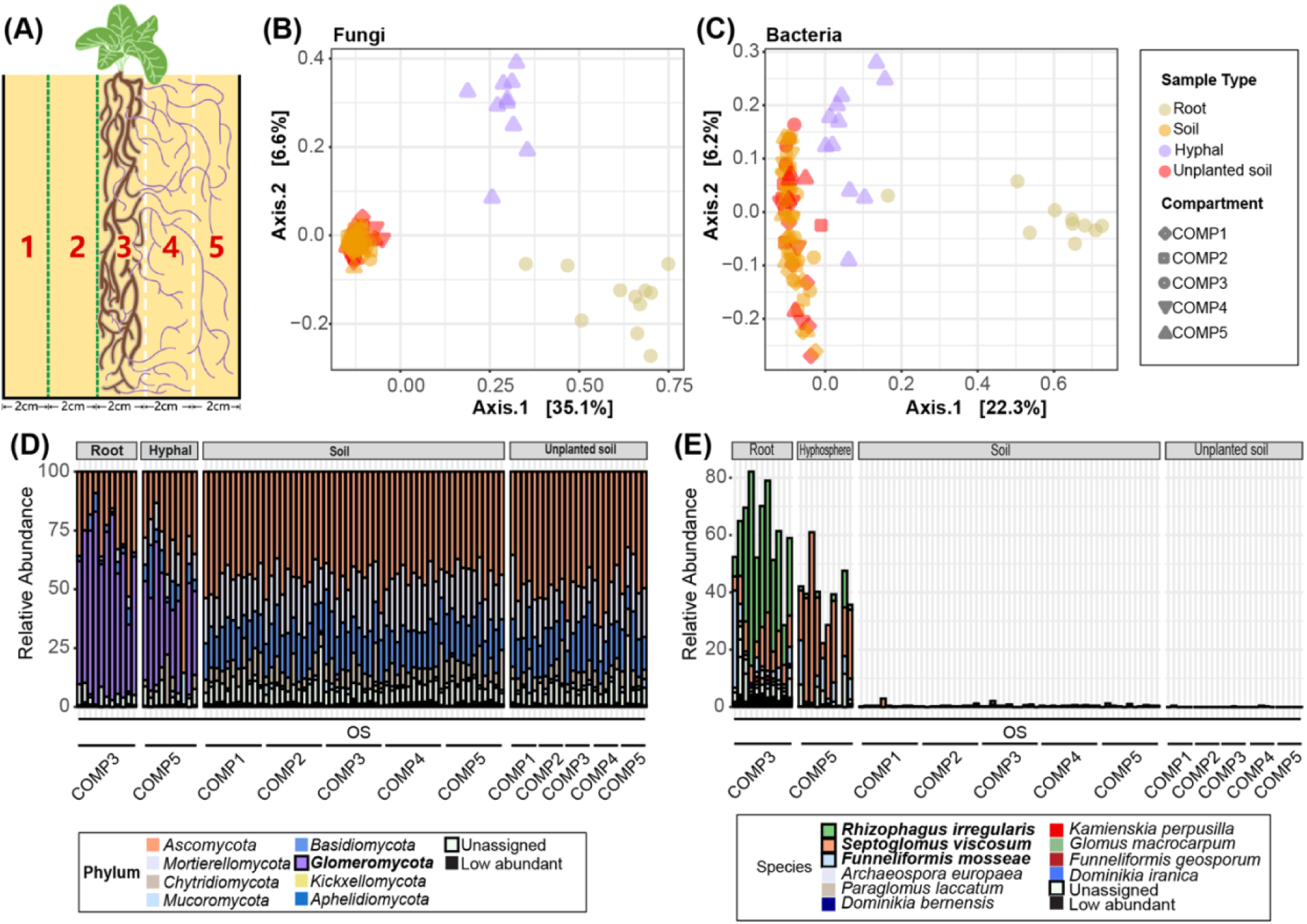
Mycorrhiza-rich hyphal samples host a bacterial microbiome that is distinct from their surrounding soil. **(A)** Schematic representation of the 5-compartment microcosm in Experiment II. All compartments were filled with 30% non-sterilized organic soil (OS), mixed with Oil-Dri and sand. Roots are contained in COMP3 by 30-µm filters (white dashed lines), whereas extraradical AM hyphae are restricted from COMP1 and 2 by 1-µm filters (green dashed line). **(B)** PCoA of fungal communities using Bray-Curtis distances in root, soil and hyphal samples of plants growing in OS. **(C)** PCoA of bacterial communities in root, soil and hyphal samples of plants growing in OS. Colors in **(B)** and **(C)** indicate different sample types. Shapes in **(B)** and **(C)** depict different compartments. **(D)** Relative abundance of fungal phyla in root (COMP3), soil (COMP1 to 5) and hyphal samples (COMP5) in Experiment II. Colors represent the distinct phyla. Phyla with relative abundance below 1% were aggregated and categorized as lowly abundant. **(E)** Relative abundance of *Glomeromycota* spp. in root, soil and hyphal samples in Experiment II. Colors represents the distinct AM fungal species.

In contrast to our expectations, we did not find a strong influence of plant growth on the soil microbiome. The soil fungal and bacterial communities of the 5 distinct compartments in the microcosms with plants were not significantly different from each other (PERMANOVA; Fungi, R^2^ = 0.077, F = 1.052, *p* = 0.257; Bacteria, R^2^ = 0.087, F = 1.095, *p* = 0.101), whereas all soil samples group together and away from the root and hyphal samples in PCoA (Fig. 2B, 2C). Nonetheless, both the bacterial and fungal communities in the root-containing COMP3 (Fig. S2) differed significantly from COMP3 soil communities of unplanted microcosms (Table S3). Moreover, the fungal community of COMP4 and the bacterial community in COMP2 were significantly affected by the presence of Prunella roots in the adjacent COMP3 and differed significantly from the same compartments in the unplanted microcosms (Table S3). This shows that roots do affect the soil microbial community of COMP3 and that root exudates can, to a lesser extent, also reach and affect the microbial communities of the adjacent COMP2 and 4. The roots however do not affect the outer COMP1 and 5. Furthermore, we were able to isolate hyphae from COMP5, and these hyphal samples are enriched with *Glomeromycota*. Moreover, the hyphal samples also contain bacterial communities that are distinct from the surrounding soil (Fig. 2C), in line with observations made in Experiment I (Figure 1C). Sample type (root, hyphal, or soil) explained 40.8 % of the variation in fungal communities and 18% of the bacterial communities over all compartments, while the presence of Prunella roots explained only 2% of the difference between unplanted and planted microcosms for fungal communities and 1.7% of the difference for bacterial communities (Table S3).

*Glomeromycota* again dominated the fungal community of both root and hyphal samples (RA of 61% and 40%, respectively; Fig. 2D). In addition to *Rhizophagus irregularis* and *Septoglomus viscosum* (the *Glomeromycota* spp. that were found abundantly in our Experiment I), we found *Funneliformis mosseae* to be also abundantly present in the root and hyphal samples of our Experiment II (Fig. 2E). Here, we found that the hyphal samples consisted of fungal and bacterial communities that were significantly different from the soil microbial communities in COMP5, which reflects the original soil from which these microbes were initially acquired (Fig. 2B, 2C, Table S4).

### Bacteria on hyphae derive from soil and root

We subsequently focused on the bacterial communities to better understand the hyphal microbiome assembly. In both Experiment I and II, we observed that the bacterial community occurring on hyphae is different from those on soil and root samples. In Experiment I, we detected a total of 5,139 bacterial amplicon sequence variants (ASVs), of which 289 ASVs occurred in root, soil as well as hyphal samples (Fig. 3A). These shared ASVs account for 33.1% of RA in hyphal samples, and 35.1% of RA in root samples, but make up only 10% of RA in soil samples. Root and soil samples uniquely share each an additional 241 and 186 bacterial ASVs with the hyphal samples, respectively. The 241 ASVs shared between roots and hyphae account for 28.6% of RA in hyphal samples, whereas they represent only 5.6% of RA in root samples. Similarly, the 186 ASVs uniquely shared between soil and hyphae represent 11.2% of RA in the hyphal samples, but only 2.2% of RA in soil samples. In total, more than 70% of RA in hyphal samples are taken up by the shared ASVs from either soil, roots or both (Fig. 3B). This suggests that most bacteria on hyphae, that were isolated from the sterilized substrate in COMP5 in Experiment I, originated from the root and soil in COMP3, and likely traveled over, within, or with the hyphae into COMP5.

**Fig. 3.**
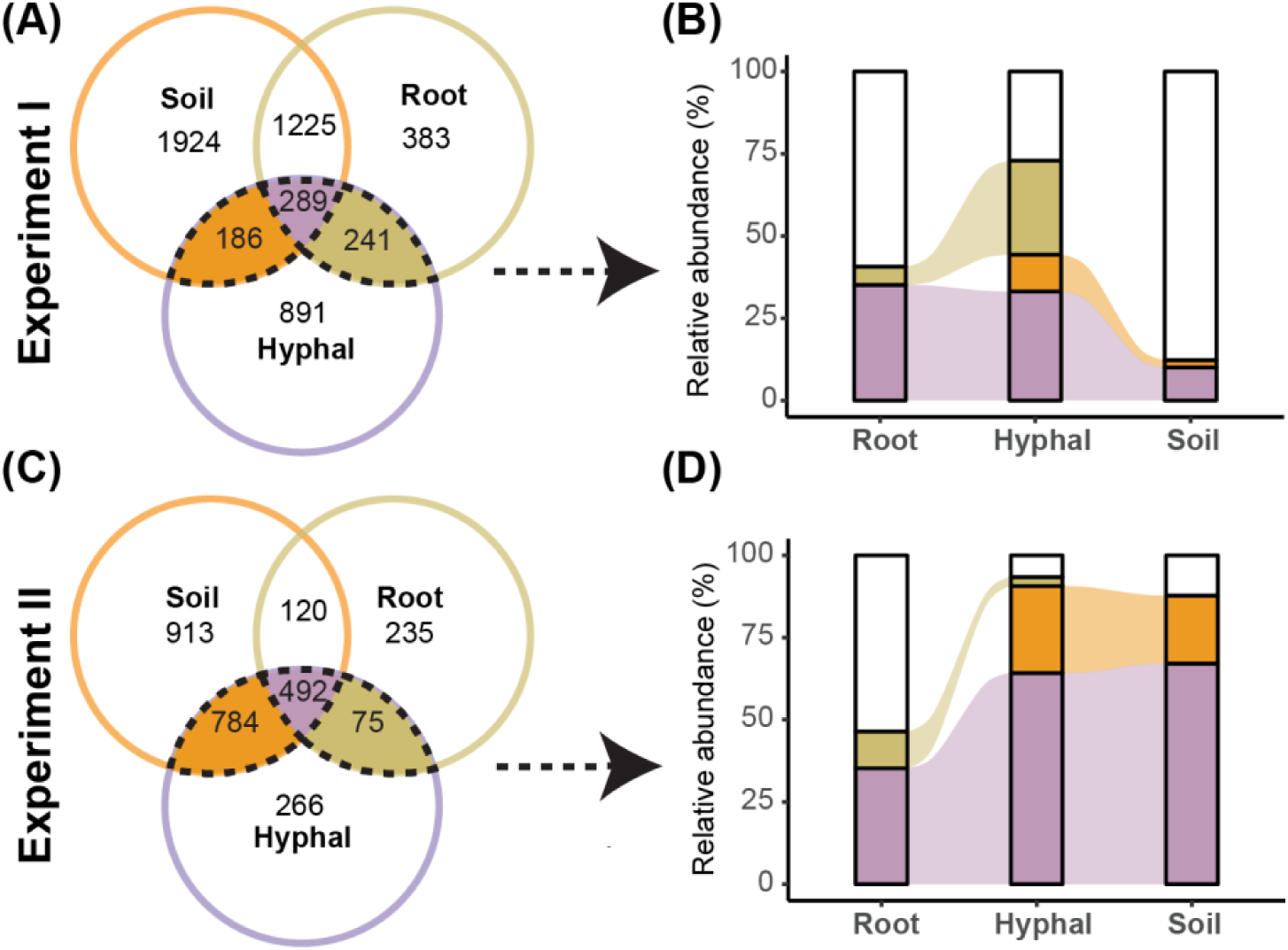
The abundance of hyphal ASVs shared with root and soil samples. **(A)** Venn diagram of unique and shared bacterial ASVs in root, hyphal, and soil samples of Experiment I. Number of ASVs are indicated for each compartment. Colors indicate bacterial ASVs shared between hyphae and soil (orange), root (green) or both (purple). **(B)** Sankey plot of hyphal samples shared ASVs’ RA in each sample types. The colors depict the hyphal ASVs either shared with soil or root or both. **(C)** Venn diagram of unique and shared ASVs in root, hyphal and soil samples of experiment II. **(D)** Sankey plot of hyphal samples shared ASVs’ RA in each sample types. Only ASVs minimum present in 3 samples are considered here.

In experiment II, however, all compartments were filled with the same non-sterilized soil substrate. Here, 492 out of a total of 2,885 bacterial ASVs were found to be shared by root, hyphal and soil samples. These ASVs account on average for 64.2% of RA in hyphal samples and 67.1% of RA in soil samples, but only 35.3% of RA in root samples. In addition, the hyphal samples uniquely share 784 ASVs with soil samples that account for 26.4% of RA in hyphal samples and 20.7% of RA in soil samples. As a result, the ASVs that together represent more than 90% of the reads in hyphal samples are also detected in soil samples (Fig. 3D). In contrast, the hyphal samples uniquely share only 75 ASVs with the root samples. That account for only 2.7% of RA in hyphal samples and 11.1% of RA in root samples. Thus, in the more natural situation of experiment II, the microbial community on hyphae is more similar to that of the surrounding soil, and only a small minority has likely travelled from the root compartment. In both cases, however, the hyphal samples constitute a microbial community that is distinct from the community observed in the soil and roots.

### Specific bacterial taxa are consistently enriched on hyphal samples

We then examined which bacterial taxa were consistently enriched in the hyphal samples to identify bacteria that strongly associate with the AM hyphae. We identified 81 bacterial genera that occurred in the hyphal samples of both experiments (Fig. 4A). These consistently present bacterial genera are more abundant in hyphal samples then soil samples, and comprise a large part of the bacterial microbiome in the hyphae of both experiments (Fig. 4B). These consistently present bacterial genera together increase from 19.9% and 16.2% in soil to 42.9% and 27.6% in the hyphal samples of Experiment I and II, respectively. Of those 81 genera, 13 genera were significantly more abundant in hyphal samples than in soil samples in both experiments (Fig. 4C), of which *Haliangium, Massillia, Pseudomonas,* genus SWB02, and *Devosia* were the most abundant. In contrast, these 13 consistently enriched bacterial genera comprise only 1.5% and 0.3% of RA in the soil samples of Experiment I and II, but represented 24.6% and 5.8% of RA in the hyphal samples of both experiments, respectively. These genera are thus consistently and specifically enriched in mycorrhiza-rich hyphal samples. Interestingly, in both experiments, *Haliangium* is by far the most abundant bacterial genus on the hyphae, taking up 6.4% and 3% of RA in Experiment I and II, respectively.

**Fig. 4.**
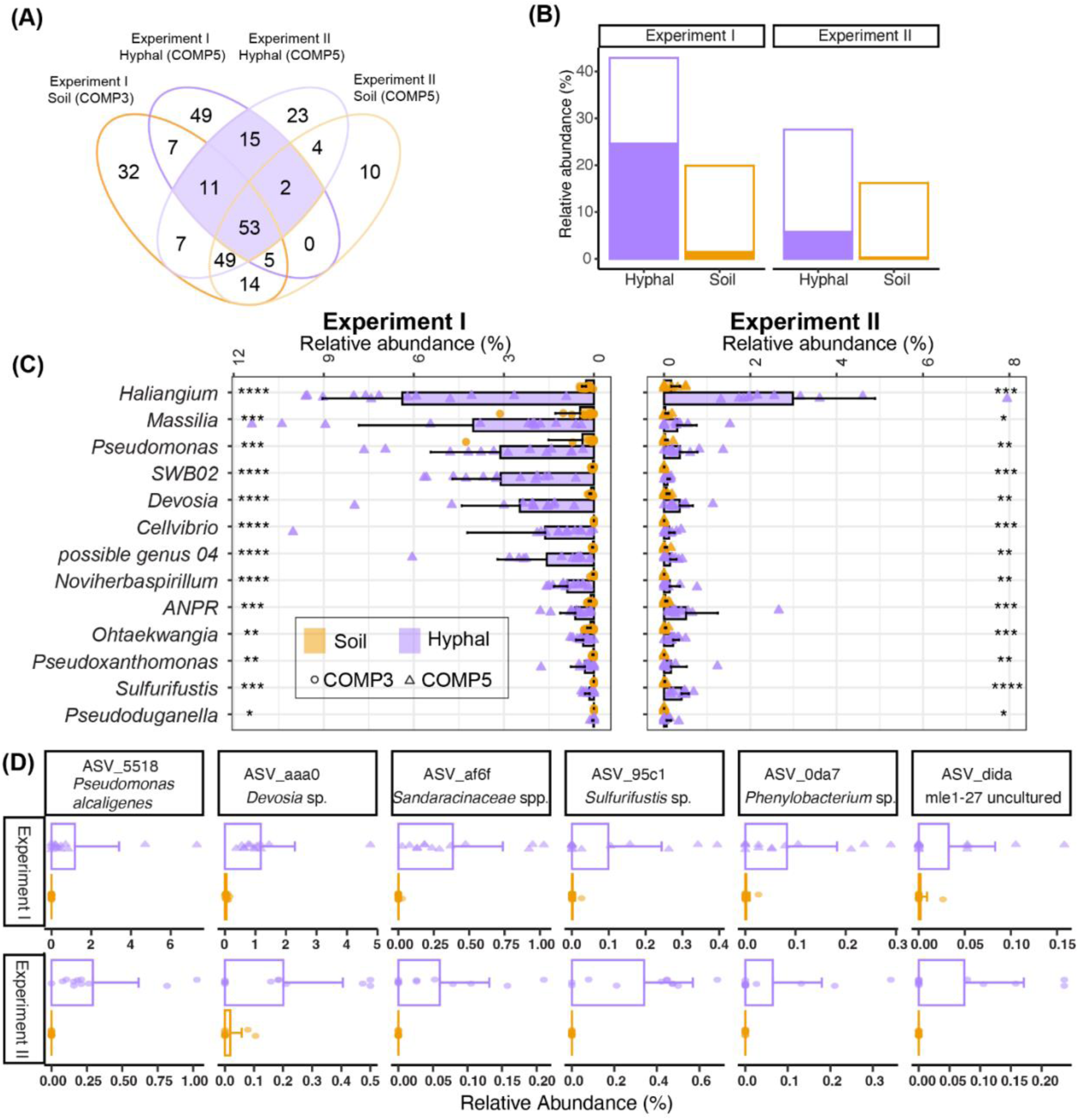
Specific bacterial genera and ASVs are consistently enriched on hyphae in both experiments. **(A)** Venn diagram showing the occurrence of bacterial genera on hyphal and soil samples across 2 experiments. Genera with relative abundance below 0.1% were aggregated and categorized as lowly abundant that are not present here. **(B)** Relative abundance of bacterial genera that are consistently occurring on hyphal samples (outline of bars) and of genera that are consistently significantly enriched in hyphal samples (filled with purple color) of Experiments I and II compared to the abundance of these same genera in soil samples (filled with orange color). **(C)** Relative abundance of genera that are consistently significantly enriched in hyphal samples across the two experiments (Wilcox-test; * *p* < 0.05, ** *p* < 0.01, *** *p* < 0.001, **** *p* < 0.0001, ns: *p* > 0.05). ANPR*: Allorhizobium-Neorhizobium-Pararhizobium-Rhizobium. **(D)** Bar plots showing the mean relative abundance of six bacterial ASVs that are consistently enriched in hyphal compared to soil samples in both Experiment I and II. Bacterial ASVs are labeled with a unique 4-letter ASV identifier and the lowest available taxonomic annotation. Colors indicate sample types; shapes of symbols indicate the microcosms of samples from which they are derived.

These results encourage us to analyze further our data at a higher taxonomic resolution. We used *Indicspecies* [37] to calculate the point-biserial correlation coefficient of an ASV that is positively associated with hyphal or soil samples. Only six bacterial ASVs were positively associated with the hyphal samples of both experiments (Fig. 4D). These ASVs are all *Proteobacteria* and belong to the genera *Pseudomonas*, *Devosia*, *Sulfurifustis*, *Phenylobacterium,* and uncultured *Myxococcales*.

In summary, certain bacterial genera appear to be consistently enriched in our hyphal samples, comprising a considerable portion of the bacterial abundance. The genus of *Halangium* represents the most strongly enriched genus and dominated the hyphal samples of our two independent experiments. Moreover, the genus *Pseudomonas* and *Devosia* stand out as they are not only consistently enriched on hyphal samples of both experiments, but each also comprises a specific ASV that is consistently associated with AM hyphae.

### Isolation of hyphosphere bacteria

To functionally characterize hyphae-associated bacteria, we isolated bacteria from mycorrhiza-rich hyphal samples collected from COMP5 in microcosms with Prunella plants of experiment I. We either placed single hyphal strands on an agar-solidified growth medium and streaked individual bacterial colonies that appeared alongside these hyphae (Fig. S5). Alternatively, we washed hyphal samples in sterile 0.9% saline water and isolated bacteria through dilution plating.

In total, we isolated 144 bacteria and determined the taxonomy of the isolates by sequencing the 16S rRNA gene (Additional file 1). The 144 isolates belong to 3 bacterial phyla and mainly represent *Actinobacteria* (72.7%), *Proteobacteria* (17.5%), and *Firmicutes* (9.8%). Of the 13 bacterial genera that were consistently enriched in hyphal samples, we isolated representatives of the genus *Pseudomonas* and *Devosia* only. Remarkably, the most abundant bacterial genus in the hyphal samples, *Haliangium,* was not represented, indicating that the *Haliangium* bacteria on the hyphae were not able to grow on the media used for isolation.

We further examined our isolate collection by matching the 16S rRNA gene of the bacterial isolates to the ASVs enriched in sequencing data of hyphal samples of the above-described experiments I and II. We isolated three *Devosia* spp. from our mycorrhiza-rich hyphal samples. These isolates have identical 16S sequences and share 99.5 % nucleotide identity with *Devosia* ASV aaa0, which was consistently enriched on hyphal samples in both experiment I and II. Interestingly, however, the isolates share 100% nucleotide identity with *Devosia* ASV e5d2, an ASV that was consistently significantly enriched on roots of Prunella plants, but not in the hyphal samples (Fig. 5A).

**Fig. 5.**
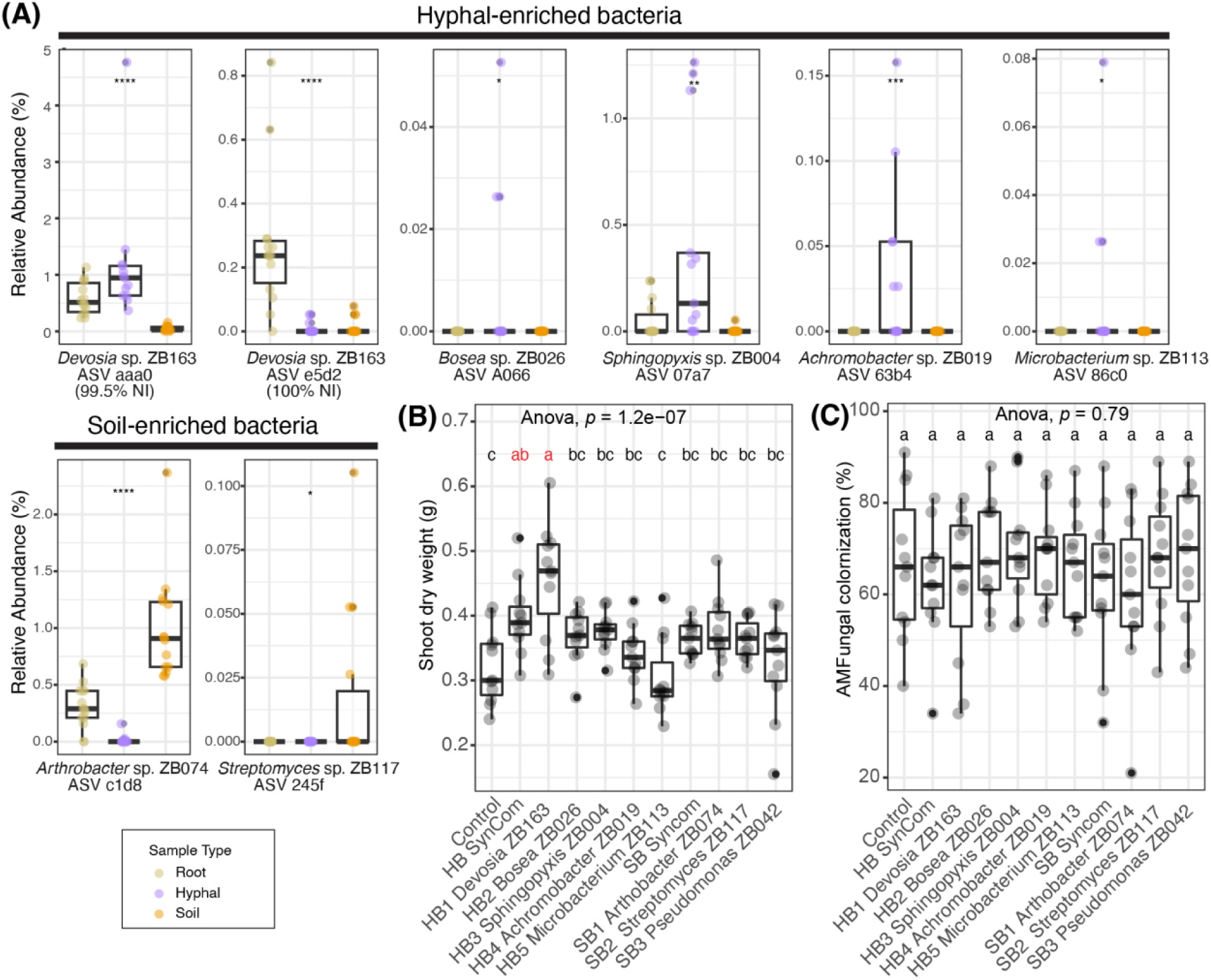
*Devosia* sp. ZB163 is isolated from fungal hyphae but thrives on the root and promotes plant growth. **(A)** Relative abundance of the selected ASVs in the root, hyphal, and soil samples in Experiment I. Sample types were indicated by color. Each selected ASVs ID was labeled together with a selected corresponding bacterial isolate with matching sequence. Significant differences are indicated with asterisk (one-way ANOVA, *p < 0.05, **p <0.01, ***p < 0.001, ****p < 0.0001). **(B)** Shoot dry weight of 9-week-old Prunella plants **(C)** AM fungi colonization percentage comparison between bacterial treatments. Significant differences of **(B) & (C)** are indicated with letters (ANOVA and Tukey’s Honest HSD test).

The 16S sequence of the single *Pseudomonas* sp. ZB042 did neither match very well with the consistently enriched *Pseudomonas* ASV 5518 (95% NI) nor any other ASV in the data set with more than 99% NI. We therefore expanded our search to identify ASVs with a shared NI of more than 99% with an ASV that was significantly enriched in hyphal samples of experiment I. In this way, we ultimately selected 5 hyphosphere bacteria (HB) from our collection of isolates that respectively represent *Devosia* ASV e5d2, *Bosea* ASV A066*, Sphingopyxis* ASV 07a7*, Achromobacter* ASV 63b4, and *Microbacterium* ASV 86c0 (Fig. 5A). These HB were subsequently used to examine their influence on the AM symbiosis. In addition, we selected 2 bacterial isolates that matched with ASVs that were enriched in soil compared to hyphal samples, and here we also included the *Pseudomonas* sp. ZB042. These soil bacteria (SB) were incorporated as control bacteria that were not associated with AM fungi.

### *Devosia* sp. ZB163 promotes plant growth in organic soil

We tested whether the selected bacterial isolates affected the symbiosis between *P. vulgaris* plants and AM fungi. To this end, we inoculated a soil-sand mixture with each of the 5 HB or the 3 SB at an initial density of 3×10^7^ CFU/g. In addition, two treatments, either combining the 5 HBs or the 3 SBs as two separate synthetic communities (HB/SB SynCom), were applied to the soil-sand mixture with a cumulative initial abundance of 3×10^7^ CFU/g. Finally, we transplanted 2-week-old prunella plants to the inoculated pots. After 9 weeks of growth in a greenhouse, we harvested the shoots of these plants and found that only plants inoculated with either *Devosia* sp. ZB163 (hereafter: *Devosia*) or the HB SynCom had significantly higher shoot dry weight than control plants (Fig. 5B). This indicates that *Devosia* can promote plant growth. All control and treatment plants in this experiment were colonized by AM fungi and the mycorrhization at the end of the experiment was not significantly affected by the distinct bacterial treatments in this experiment (Fig. 5C).

### *Devosia* sp. ZB163 promotes plant growth and mycorrhization

To explore whether plant growth promotion by *Devosia* sp. ZB163 relies on the presence of AM fungi, we depleted the indigenous microbiome by autoclaving the soil-sand mixture and again inoculated *Devosia* at an initial density of 3×10^7^ CFU/g soil prior to transplantation of *Prunella* seedlings (hereafter: *Devosia* treatment). Subsequently, 100 monoxenic *R. irregularis* spores were injected near the seedling’s roots (hereafter: AM treatment). To ensure nutrient-poor conditions and stimulate AM fungi colonization, the plants in this experiment were not provided with nutrients in addition to what was present in the soil-sand mixture.

After 8 weeks of growth under controlled conditions in a climate chamber, plants inoculated with *Devosia* had a significantly higher shoot and root weight (Fig 6A, 6B), indicating that, even without AM fungi, *Devosia* sp. ZB163 can promote plant growth. Four out of the eleven plants that were inoculated with AM fungi were bigger than control plants and the leaves of these plants were more bright green (Fig. 6F). These four plants were the only plants in which mycorrhiza had colonized the roots and, likely as a result of the mycorrhiza incidence, the average weight of roots and shoots was not affected by the AM treatment. However, plants that had been inoculated with the combination of AM and *Devosia* did have significantly higher shoot and root weights compared to the controls without AM and *Devosia*. Remarkably, 10 out of 11 plants that had received the combination of *Devosia* and AM were bright green and were colonized by mycorrhiza. This suggests that *Devosia* sp. ZB163 not only promoted plant growth directly but also improved AM establishment in this experiment. As *Devosia neptuniae* has previously been reported to fix N [38] and AM fungi are known to provide plants with both N and P [39], we measured leaf N and P content. We found that the leaves of all plants that were colonized by AM fungi contained more P (Fig 6E), while the plants that were inoculated with *Devosia* had higher N content (Fig. 6D). This suggests that *Devosia* and AM promote plant growth by stimulating the uptake of respectively N and P in a complementary manner. We hypothesized that this did not result in even higher plant growth in the combination treatment as other mineral components of the nutrient-poor soil/sand mixture also constrained the growth of plants in these experiments.

**Fig. 6.**
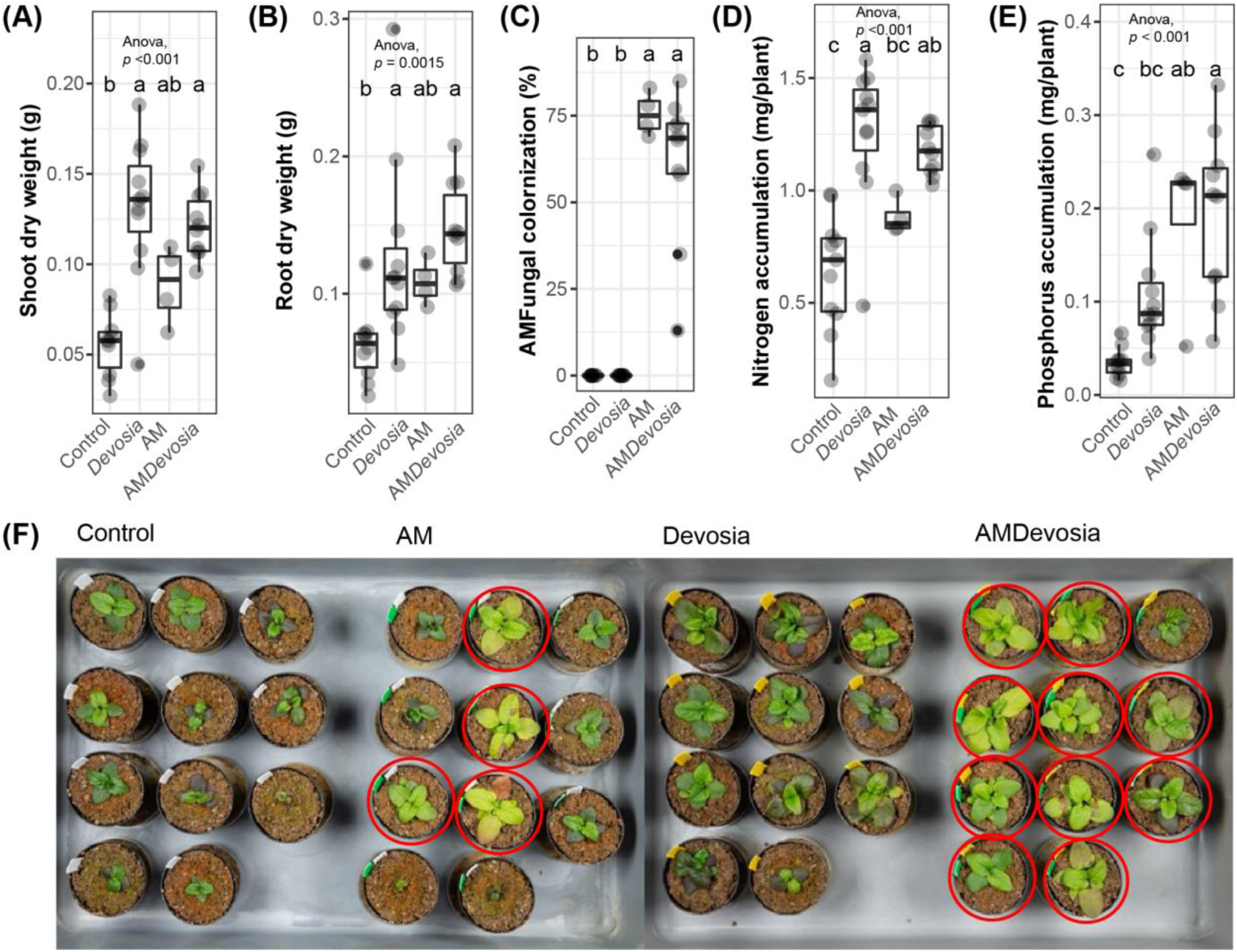
*Devosia* promotes plant growth, mycorrhization, and N accumulation. Boxplots show **(A)** shoot dry weight, **(B)** root dry weight, **(C)** percentage of each root system colonized by AM fungi, **(D)** shoot N accumulation, and **(E)** shoot P accumulation of 8-week-old Prunella plants cultivated in autoclaved soil (Control) or inoculated with *Devosia* sp. ZB163 (Devosia), *R. irregularis* (AM), or both symbionts. In the 6^th^, 7^th^ and 8^th^ week, plants were watered with modified Hoagland solution without N and P. Significant differences are indicated with letters (ANOVA and Tukey’s Honest HSD test). **(F)** Photographs of the Prunella plants immediately before harvest. Red circles indicate plants that were later found to be colonized by AM fungi.

### *Devosia* sp. ZB163 and AM fungi synergistically promote plant growth

We subsequently repeated this experiment but now provided the plants with a modified Hoagland solution that included most micronutrients but was deficient in N and P (Table S5). Again, *Devosia* promoted plant growth, but in this experiment also AM led to a significantly higher dry weight of both shoots and roots (Fig. 7A, 7B). In this experiment, AM fungi established successfully in the roots of all plants to which they were inoculated, but the mycorrhizal colonization was higher on plants that were also inoculated with *Devosia* (Fig. 7C). Notably, this combination treatment of AM and *Devosia* resulted in the significantly highest plant shoot weight among all treatments, showing that AM fungi and the *Devosia* ZB163 can synergistically promote plant growth (Fig. 7A). In line with this, we found that accumulation of N was significantly increased in plants inoculated with *Devosia* (Fig. 7D). Moreover, although accumulation of P increased in plant inoculated with AM only, the plants inoculated with both AM and *Devosia* accumulated significantly more N and P (Fig. 7E).

**Fig. 7.**
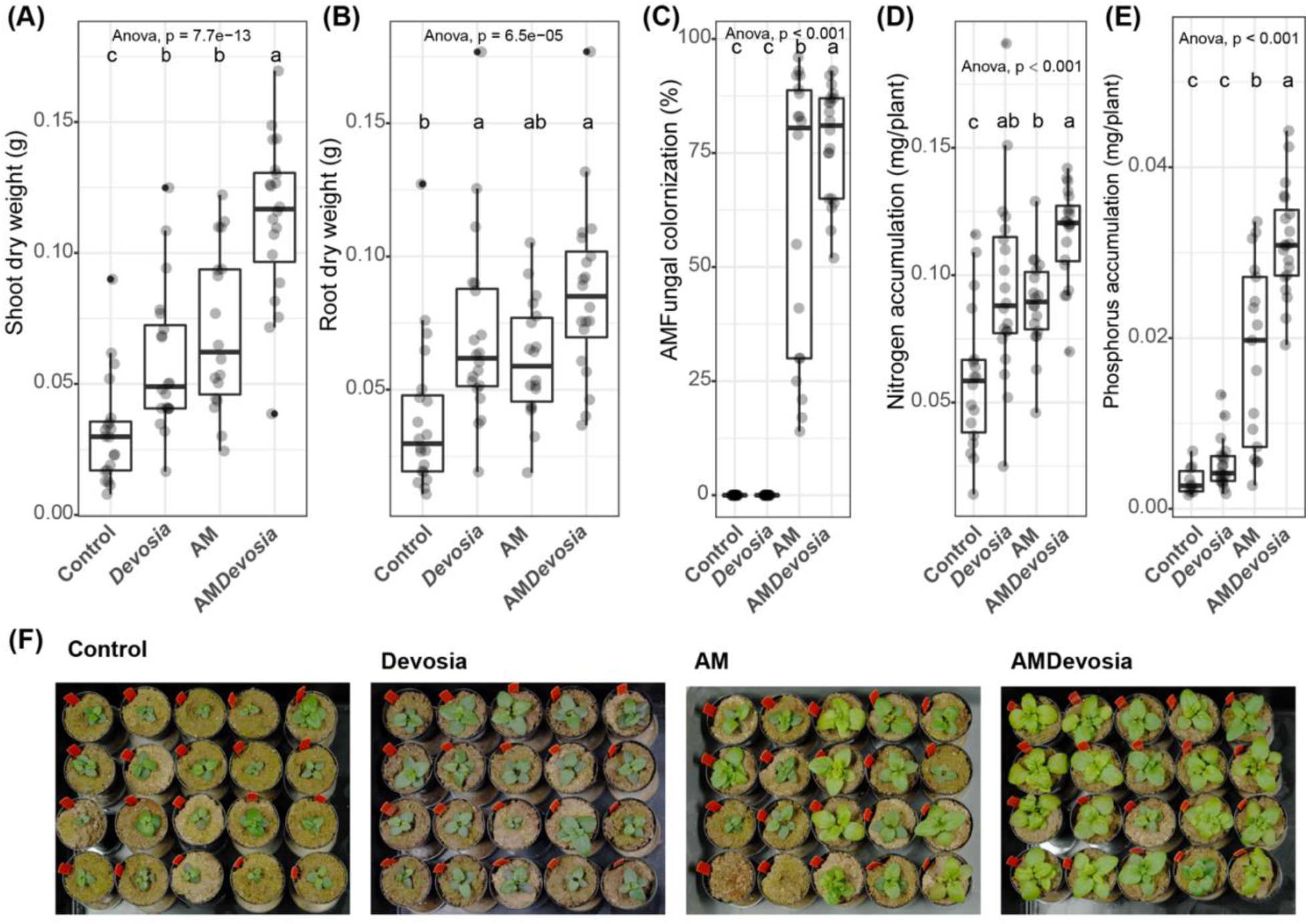
*Devosia* sp. ZB163 and AM fungi can synergistically promote plant growth and plant N and P accumulation. Boxplots show **(A)** shoot dry weight, **(B)** root dry weight, **(C)** percentage of each root system colonized by AM fungi, **(D)** shoot N accumulation, or **(E)** shoot P accumulation of 8-week-old Prunella plants cultivated in autoclaved soil (Control) or inoculated with *Devosia* sp. ZB163 (Devosia), *R. irregularis* (AM), or both symbionts. Plants were regularly watered with modified Hoagland solution deficient in a source of N and P. Significance differences are indicated with letters (ANOVA and Tukey’s Honest HSD test). **(F)** Photographs of the Prunella plants immediately before harvest. Two AM-treated plants died shortly after transplantation and were not considered in panels **A**-**E**.

We subsequently quantified the absolute abundance of *Devosia* by sequencing 16S rRNA gene amplicons of DNA isolated from the roots of plants used in this experiment and spiked with a known amount of 14ng DNA [40]. We detected low amounts of *Devosia* on the roots of plants that were not inoculated with *Devosia*, indicating that some level of cross contamination occurred in our experiment (Fig. 8A). Nonetheless the numbers of *Devosia* were significantly higher on roots that were inoculated with *Devosia*.

**Fig. 8.**
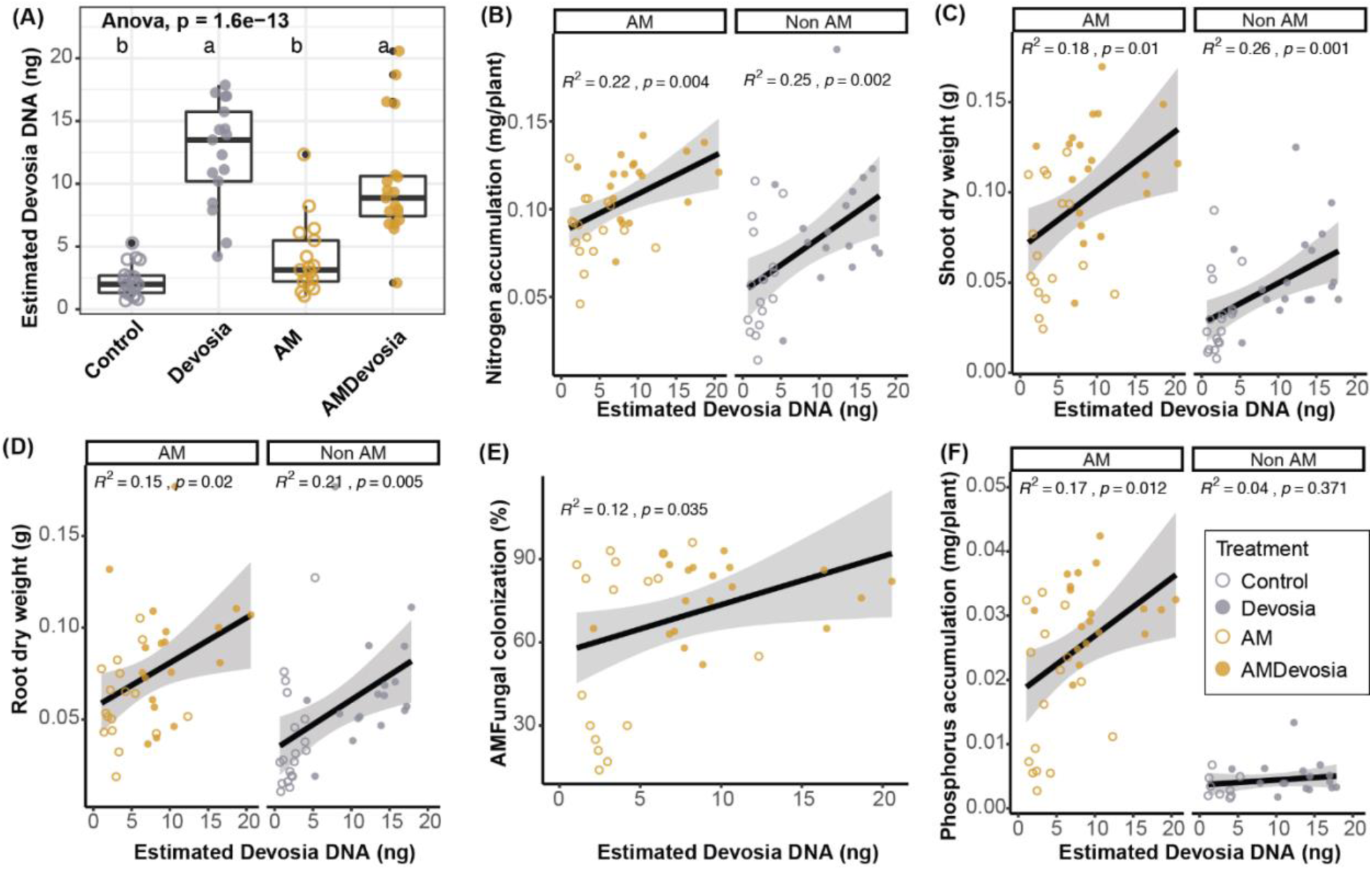
Abundance of *Devosia* sp. ZB163 significantly correlates with plant weight, mycorrhization, and N and P accumulation. **(A)** Boxplot of the absolute abundance of *Devosia* DNA on roots of plants in sterilized soil inoculated with a mock solution (Control), *Devosia* sp. ZB163 (*Devosia*), *R. irregularis* (AM), or both symbionts. Letters indicate significant differences as determined by ANOVA with Tukey’s HSD test. **(B-E)** Scatter plots of the correlation between the absolute abundance of *Devosia* DNA and **(B)** total plant N accumulation, **(C)** shoot dry weight, **(D)** root dry weight **(E)** hyphal colonization, and **(F)** total plant P accumulation. Correlations and probabilities thereof are determined using linear regression.

We subsequently analyzed the correlation between absolute *Devosia* abundance and several parameters. We observed that, independent of AM presence, *Devosia* abundance positively correlates with plant N accumulation (Fig. 8B), but also with shoot and root dry weight (Fig. 8C, 8D). This, together with the observed causal effects, shows that *Devosia* sp. ZB163 can directly stimulate plant growth and N uptake. Moreover, the absolute abundance of *Devosia* significantly correlates with the percentage of AM fungi colonization (Fig. 8E), suggesting further that *Devosia* indeed accelerates the colonization of plant roots by AM fungi. In line with this, we observed that *Devosia* abundance correlates significantly with increased P accumulation, but only in presence of AM (Fig. 8F), and that the percentage of root length colonized by AM hyphae correlates with P accumulation (Fig. S6). Together, these data show that *Devosia* can stimulate plant growth directly, likely by increasing N uptake, but also indirectly by promoting AM fungi colonization and corresponding P uptake.

### *Devosia* sp. ZB163 lacks genes required for atmospheric N fixation

The genome of *Devosia* sp. ZB163 was subsequently sequenced using the Illumina Novoseq platform (Génome Québec, Canada) resulting in a sequenced genome of approximately 4.6 Mb that was predicted to have 4486 coding sequences (CDSs) and a GC content of 65.7%. As we found that *Devosia* sp. ZB163 promotes plant N uptake, we subsequently performed a reciprocal BLASTp to search for orthologues of known N-related genes (Table S6). We first explored the *Devosia* genome for genes that are required for atmospheric N fixation. The *nifADHK* gene cluster typically encodes the molybdenum nitrogenase complex that is most commonly found in diazotrophs (Dixon & Kahn, 2004). However, we found orthologues of neither *nifA, nifD, nifH* nor *nifK* in the genome of ZB163 using translated amino acid sequence of these genes from *Devosia neptuniae*, *Sinorhizobium meliloti*, *Bradyrhizobium japonicum* and *Klebsiella pneumoniae* [38, 41-43]. Next, we blasted the *Devosia* sp. ZB163 genome to a *nifH* database that contains 34420 *nifH* sequences, but again did not find a hit for *nifH* in the genome of ZB163. Finally, also the gene clusters *vnfHDGK* and *anfHDGK* encoding the less common nitrogenase complexes were not detected in the *Devosia* sp. ZB163 genome [44]. This strongly suggests that unlike other *Devosia* isolates, *Devosia* sp. ZB163 is not able to fixate atmospheric N.

However, bacteria can also increase the amount of N that is available to plants through the mineralization of organic N. The ammonification process in the soil mineralizes organic N to ammonia and the organic soil used in this study was previously reported to slowly-release urea [45]. Urea, as an organic N source, is subsequently catalyzed by urease to ammonia that can be subsequently supplied to plants. Using protein sequence from *Devosia rhizoryzae*, *Devosia oryziradicis* [46], we detected the presence of the gene clusters *UreDFG* and *UrtABCDE* that are required to catalyze the hydrolysis of urea, forming ammonia and carbon dioxide. Besides ammonia, plants can also take up nitrate. Nitrification bacteria catalyze ammonium to nitrate with amoA gene. Again, we did not detect any amoA orthologs in the *Devosia* genome using the translated amino acid sequences of these genes from *Nitrosomonas europaea* [47].

## Discussion

Plant root microbiomes are known to play important roles in plant growth and plant health [11]. Here we investigated whether AM fungi, that are part of the plant root microbiome, are themselves also similarly able to interact with microbes. AM fungi do not only transfer mineral nutrients to the host plants, but also relocate 5-20% of photosynthates from the plant to the surrounding environment [48, 49]. As such, the AM hyphae provide space and nutrients for microbes to grow on and has been shown that the AM hyphosphere microbiome is different from the bulk soil [22, 23]. While some studies assessed bacterial communities associating with AM hyphae, so far, no studied isolated bacteria from AM hyphae and test the impact on plant growth and mycorrhization. To resolve this gap of knowledge, we conducted experiments in compartmentalized microcosms, and we sampled hyphae that grew from a compartment with plant roots into the outer compartment of the microcosms, from which roots were restricted. These hyphal samples were strongly enriched in *Glomeromycota*, the division of the obligate biotrophic fungi that form arbuscular mycorrhiza. Moreover, we were unable to isolate these hyphae from the same compartment of unplanted microcosms, which demonstrates that a large part of these hyphae is likely formed by extraradical hyphae of obligate fungal biotrophs that extend from the prunella roots in these microcosms. Nonetheless, although most bacterial isolates were likely isolated from AM fungi, it is possible that some were isolated from other fungi (e.g., *Chitriodiomycota* were also common in some microcosms).

We found that the bacterial communities in our hyphal samples are distinct from the surrounding soil. Although a select set of microbes appear to have traveled from the root compartment to the hyphal compartment, the majority of the microbes on hyphae are shared with the surrounding soil but changed in abundance on the hyphae. AM hyphae thus selectively assemble a bacterial hyphosphere microbiome and this confirms other studies [22, 24, 25, 50]. In our first two experiments, *Haliangium* is the most abundant bacterial genus in our hyphal samples. Representatives of this genus have previously been isolated from soil samples and, as bacterivore *Haliangium* spp. have been found to prey on bacterial species, it has been hypothesized that *Haliangium* spp. shape the soil microbiome through bacterivory [51-54]. The abundance of *Halangium* spp. on AM-fungi-rich hyphae suggests they are important for AM fungi and hyphosphere communities. Unfortunately, we were unable to isolate *Halangium* spp. from AM-fungi-rich hyphae in this study using the conventional growth media, perhaps because these *Halangium* spp. are bacterivores that obtain energy and nutrients entirely from the consumption of bacteria. It will be interesting to explore their role in the AM hyphosphere in the future.

In addition to *Haliangium*, also the genera *Pseudomonas* and *Devosia* were consistently enriched in the hyphal samples of our experiments. Previously, *Pseudomonas* strains have been identified as mycorrhiza helper bacteria that promote the colonization of both ectomycorrhizae and arbuscular mycorrhizae in multiple studies [25, 27, 55, 56]. A recent study even suggested that the recruitment of *Pseudomonas* strains reduces N_2_O emissions from soil [25]. Our results suggest that the beneficial effect of *Pseudomonas* bacteria on AM fungi is reciprocated by the AM fungi, who can also specifically promote the growth *Pseudomonas* spp.

*Devosia* spp. have not previously been found in association with AM fungi, but we found this genus to be consistently enriched in mycorrhiza-rich hyphal samples. We were able to isolate *Devosia* sp. ZB163 from the mycorrhiza-rich hyphal sample, but the 16S rRNA gene sequence *Devosia* sp. ZB163 was a perfect match to a *Devosia* ASV that was especially abundant in root samples. Although this might suggests that *Devosia* sp. ZB163 operates largely on the roots of Prunella plants, *Devosia* sp. ZB163 is nonetheless also present on hyphal samples. As fungal hyphae are recognized as highways of bacterial movement [57], it will be interesting to investigate the role of mycorrhizal hyphae in transport of this bacterium to new hosts. Fungus-mediated transport of *Devosia* sp. ZB163 would benefit this bacterium, the fungus that transports it, as well as their mutual host plant. On prunella roots, *Devosia* sp. ZB163 can stimulate plant growth directly, but it also enhances the mycorrhizal colonization process and thus functions as a mycorrhization helper bacterium [27].

*Devosia* sp. ZB163 also promotes the uptake of N by the plant as evidenced by the increased amount of total N in Prunella plants that were inoculated with the isolate. To have insight into the mechanism by which *Devosia* sp. ZB163 promotes N uptake by Prunella, we sequenced the genome of *Devosia* sp. ZB163 and searched for genes involved in N conversion. Whereas our analysis suggests *Devosia* sp. ZB163 is not involved in N fixation or nitrification, we did identify gene clusters that are putatively used for the decomposition of urea, which is a critical process for ammonification in soil [58] and which could improve plant N availability [59].

Although AM fungi require considerable amounts of N for their own development, they can still contribute to the N uptake by the host plant [60]. AM fungi take up inorganic N outside the roots, mostly as ammonium [61, 62], incorporate it as glutamine, translocate the N from the extraradical to the intraradical mycelium as arginine, and once inside the root cells, convert the arginine into urea, from where the N is finally transferred as ammonium to the host [5]. Hence, urea is an important precursor of ammonium [61], and it is tempting to speculate *Devosia* sp. ZB163 also operates as an endosymbiont, as observed for other AM-associated bacteria [63], and facilitates transfer of inorganic N to the host plant inside the intraradical hyphae by converting urea into ammonium. Consistent with this, our co-inoculation with *Devosia* sp. ZB163 and AM fungi in Prunella plants increased mycorrhization, suggesting a bacterial ability to enhance AM fungi growth, and also led to the highest accumulation of N in the host plant. Future research should attempt to characterize whether *Devosia* sp. ZB163 can operate as an endosymbiont of AM fungi.

Alternatively, *Devosia* sp. ZB163 might induce a response in the plant that enhances N uptake. For example, an *Achromobacter* sp. in the root of oilseed was found to stimulate the uptake rate of nitrate by stimulating the plant’s ionic transport system while simultaneously promoting the formation and length of root hairs[64]. It will be intriguing to find out whether *Devosia* sp. ZB163 similarly promotes the formation of an extensive root system in Prunella plants, as extensive root branching likely also affects the rate of mycorrhization [27]. In line with this hypothesis, we did see a significant correlation between root dry weight and the abundance of *Devosia* sp. ZB163 on the roots in our experiments.

*Devosia* sp. ZB163 by itself did not affect plant P content, but in presence of the mycorrhiza, the abundance of *Devosia* sp. ZB163 was significantly correlated with increased P accumulation. This shows that, although *Devosia* sp. ZB163 does not itself provide P to the plant, it can indirectly provide extra P by stimulating mycorrhization and/or the mycorrhizal functioning. In line with this, we found that the combined treatment of AM fungi and *Devosia* sp. ZB163 can lead to more growth promotion than either microbe alone.

### Conclusions

Overall, our study reveals that the microbiome of AM-fungi-rich hyphal samples is distinct from the surrounding soil and that specific bacteria are selected on fungal hyphae. We found that *Halangium*, *Pseudomonas*, and *Devosia* were consistently enriched in our hyphal samples. *Devosia* sp. ZB163 acts as a mycorrhization helper bacterium, promoting the mycorrhization of Prunella plants and indirectly providing extra P to the plant. The combination of AM fungi and *Devosia* sp. ZB163 results in more growth promotion than either microbe alone. These results provide new insights into the importance of the AM fungal microbiome and highlight the potential of beneficial bacteria such as *Devosia* for improving plant growth, nutrition, and health. Further studies are needed to explore the role of these bacteria in the AM fungal hyphosphere. Mycorrhizae are a long-standing promise for sustainable agriculture and their successful application could reduce the requirements of crop fertilizers. Our study suggests that the performance of mycorrhiza and crops in the agricultural field might benefit considerably from the application of mycorrhiza helper bacteria, such as *Devosia* sp. ZB163.

## Methods

### Soil collection

The organic soil (OS) and conventional soil (CS) used in this study were derived from the Farming System and Tillage experiment (FAST) site [35]. The FAST site was established in 2009 near Zürich (latitude 47°26’ N, longitude 8°31’ E) and the plots in this field have since undergone either conventional or organic management. The soil was collected in April 2019 and March 2020 for experiment I and II respectively. The top layer of vegetation (2 cm) was removed, and a 30 cm depth of soil was excavated from the field. The soil was passed through a 2 mm sieve and stored at 4 ℃ before use.

### Description of microcosms and plant growth conditions

#### Experiment I

Microcosms were constructed of 20×10×19 cm (L×W×H) that were divided into 5 equal compartments (Fig. 1A). The compartments were separated from each other by 30-μm nylon filters that allows hyphae to pass through but not roots. COMP1 and COMP2 were separated by a 1-μm filter that also blocked hyphae. The middle compartment (COMP3) was filled with 1200 g of a mixture of 30% non-autoclaved soil (either OS or CS), 4% autoclaved Oil-Dri (Damolin GmbH, Oberhausen, Germany), and 66% autoclaved sand. This compartment acted as soil inoculum. The outer compartments (COMP1, COMP2, COMP4, and COMP5, respectively) were each filled with 1200g of sterilized outer substrate (8% autoclaved soil (either OS or CS), 6% autoclaved Oil-Dri and 86% autoclaved sand). All autoclaved substrates used in this study were heated to 121℃ for 45 mins twice. Seven replicate microcosms were set up for OS and CS, respectively.

*Prunella vulgaris* (henceforth Prunella) seeds were vapor-phase sterilized by exposure to chlorine gas for 4 hrs. To this end, chlorine gas was generated by adding 3.2 ml 37% HCl to 100 ml Bleach (Hijman Schoonmaakartikelen BV, Amsterdam, NL). The seeds were sown on half-strength Murashige and Skoog basal agar-solidified medium (Sigma Aldrich, St. Louis, MO, USA). The plates with seeds were subsequently incubated in a climate chamber (Sanyo MLR-352H; Panasonic, Osaka, Japan) under controlled conditions (light 24℃, 16 h; dark 16℃, 8 h). Seven two-week-old seedlings with roots of approximately ∼0.5 cm length were transplanted to the middle compartment of the microcosms. The plants in the microcosms were allowed to grow in the greenhouse (Reckenholze, Agroscope, Zürich, CH) with a 16h photoperiod at 24℃ alternated with 8h of darkness at 16℃. Plants were watered with 120 ml H_2_O 2-3 times per week.

#### Experiment II

To investigate the effect of an actively growing AM mycelium on the indigenous soil microbiome, we filled each of the compartments of the microcosm described above with 750 g of a mixture of 30% non-autoclaved OS, 4% autoclaved Oil-Dri (Damolin GmbH, Oberhausen, Germany) and 66% autoclaved sand. In this experiment, COMP1 and COMP2, and COMP2 and COMP3 were separated by 1-μm nylon filters to generate two AM-fungi-free compartments. COMP3 and COMP4, and COMP4 and COMP5 were separated by 30-μm nylon filters to create 2 compartments that could be colonized by extraradical AM hyphae (Fig. 3a). We set up 11 biological replicates with Prunella plants in the center compartment (as described above) and 5 biological replicates of unplanted control. The plant growth conditions were similar to those described above for Experiment I, but the experiment was executed in a greenhouse at the botanical gardens of Utrecht University.

### Harvest and mycorrhizal root colonization analysis

In both experiments, the shoots of 3-month-old plants were cut at the soil surface, dried at 70℃ for 48h, and weighed. The microcosm soil was sampled by deconstructing the microcosm compartment by compartment, homogenizing the soil of each compartment, and collecting approximately 500 mg of soil in 2-ml tubes. For sampling of AM hyphae, 30 g of soil substrate was collected from COMP5 and stored in a 50-ml tube at -20℃. The plant roots in COMP3 were collected by carefully removing soil from the roots and rinsing them under the running tap. For each microcosm, a 1-cm-long fragment of the rinsed root was cut weighed and stored in 50% ethanol for mycorrhizal root colonization analysis. Another 1-cm-long fragment of roots was cut, weighed, and stored at -80℃ for root microbiome analysis. The rest of the roots were weighed, dried at 70℃ for 48h and weighed again. From this root, water content was determined and the total root dry weight was calculated based on the combined fresh weight of all three root samples.

To check the mycorrhizal colonization of the roots, the root fragments stored in 50% ethanol were cleared in 10% KOH and stained with 5% ink-vinegar following a protocol described by Vierheilig *et al.* [65]. The percentage of total mycorrhiza colonization and frequency of hyphae, arbuscules, and vesicles were scored following the magnified intersections method by checking 100 intercections per sample at the microscope using a 200x magnification [66].

### Sampling of fungal hyphae from soil substrate

To sample fungal hyphae, we modified a wet sieving protocol typically used to collect mycorrhiza spores [67]. The schematic graph of the fungal hyphae extraction procedure is shown in Fig. S7. Briefly, 500μm, 250μm, and 36μm sieves were surface sterilized to minimize irrelevant environmental microbes present in a hyphal sample by submersing in 0.5% sodium hypochlorite for 20 mins, then submersed in 70% Ethanol for 10 mins [68]. The sieves were stacked together with the biggest filter size on top and the smallest filter size at the bottom. 25 g of soil substrate from COMP5 was placed on the top sieve. The small particles were washed down, and soil aggregates were broken down with sterilized water. The leftovers on all sieves were washed off into Petri dishes. Then, approximately 0.1 mg hyphae were picked from the samples in the Petri dishes using a set of flame-sterilized tweezers under a binocular microscope. We concentrated the hyphae in a single 1.5-ml tube filled with 0.2ml 30% glycerin per compartment. This was then considered a hyphal sample (Fig. S7, S8). The hyphal samples were stored at -80℃ until DNA extraction.

### Soil, root, and hyphal microbiome profiling

For Experiment I, the soil and root samples from COMP3 and concentrated hyphae samples from COMP5 were characterized by conducting 16S and ITS amplicon sequencing. For Experiment II, the soil samples (both planted and unplanted soil) from COMP1, 2, 3, 4, and 5, root samples from COMP3, and concentrated hyphae samples from COMP5 were characterized by conducting 16S and ITS amplicon sequencing. DNA extraction from soil, root, and hyphal samples was performed using the DNeasy PowerLyzer PowerSoil Kit (Qiagen, Hilden, Germany). The root and soil samples were homogenized in the kit’s PowerBead solution for 10 mins at 30 m/s twice using a Tissuelyser II. The hyphal samples were homogenized in PowerBead solution for 2 mins at 30 m/s 4 times with the Tissuelyser II. The rest of the DNA extraction steps followed the manufacturer’s instructions. Extracted DNA was quantified using Qubit dsDNA BR Assay Kit and Qubit Flex Fluorometer (Thermo Fisher Scientific, Waltham, MA, USA).

DNA was amplified following a two-step PCR protocol. In the first step, we amplified bacterial 16S rRNA gene V3-V4 region (341F and 806R) [69], fungal ITS2 (5.8SFun and ITS4Fun) [70, 71] using primers described in Table S7. The microbial communities were amplified in 24µl reaction volume containing 7.5 ng DNA template, 12 µl KAPA HiFi HotStart ReadyMix (F. Hoffmann-La Roche AG, Basel, Switzerland), 2.5 µl 2 µM (bacterial and fungal) forward and reverse primers and the rest volume were supplemented by MilliQ-purified water. The resulting PCR products were purified using AMPure XP beads (Beckman Coulter, High Wycombe, UK) according to the manufacturer’s instructions. The purified PCR products were then used as template DNA in the second PCR. The second PCR was performed similarly to the aforementioned but using primers from the Illumina Nextera Index Kit v2 that contain an error-tolerant 6-mer barcode to allow multiplexed library sequencing. The resulting PCR products were then cleaned-up again using AMPure XP beads. The two-step PCR were processed on a thermocycler (Hybaid, Ashford, UK) with cycling conditions as described in Table S8. The cleaned-up PCR products were quantified using Qubit dsDNA BR Assay Kit and Qubit Flex Fluorometer. Equal amounts of PCR product (2 µl 4nM) were pooled and sequenced on an Illumina MiSeq Sequencer (Illumina, San Diego, USA) using a paired-end 300bp V3 kit at Utrecht Sequencing Facility (www.useq.nl).

### Isolation of hyphae-adhering bacteria

In Experiment I, we sampled hyphae from microcosms with *Prunella vulgaris* (henceforth *Prunella*) plants. Here, we used two strategies to isolate AM associated bacteria from those hyphal samples. The first strategy was to place hyphae on agar plates directly and let the bacteria attached to the hyphae grow. Briefly, concentrated hyphal samples stored in -80 ℃ were thawed at room temperature. In a sterile laminar flow cabinet, the hyphae were gently rinsed in a sterile 3.5% Na_4_P_2_O_7_ solution to disaggregate small soil particles [20], then rinsed twice with sterile 0.9% saline water in a 2-ml tube, and subsequently transferred to a sterile petri-dish with sterile saline water. From there, single hyphal strands were picked from the saline water onto an agar plate using sterile tweezers. A maximum of eight hyphae were placed evenly distributed on a single agar plate (Fig. S5 A, B, C, D).

The second strategy was to suspend hypha-adhering bacteria in solutions and culture serial diluted solutions on agar plates. Briefly, the hyphae were concentrated, gently rinsed by a sterile 3.5% Na_4_P_2_O_7_ solution and saline water as described above. Rinsed hyphal samples were transferred to 900µl sterile 0.9% saline water, followed by rigorous shaking for 40s at 5.5 m/s in a Tissuelyser II (Qiagen, Hilden, Germany). Serial dilutions of these samples were then plated on agar-solidified culture media (Fig. S5 E, F). In both of the above strategies, seven distinct agar-solidified media were used to culture hyphae-adhering bacteria (Table S9). Single bacterial colonies were picked after 3-21 days of incubation at 28 ℃ and streaked on ISP2 agar medium (Yeast extract, 4g/l; Malt extract, 10g/l; Dextrose, 4g/l; Agar, 20g/l; pH =7.2). After 3-7 days of incubation at 28℃, isolates were examined for purity, and overnight cultures of single colonies in medium at 28℃ were stored in 25% glycerol at -80℃ for future use.

### Characterization of bacterial isolates and mapping to ASVs

To characterize the bacterial isolates, we used a pipette tip to transfer a single colony growing on ISP2 medium to 50µl of sterile water. The bacterial suspension was then incubated at 95℃ for 15mins and immediately cooled on ice. Subsequently, the bacterial lysate was centrifuged at 10,000×g for 1min to remove cell debris. Two microliters of supernatant were taken as DNA template to amplify the 16S rRNA gene using 2.5µl 27F and 2.5µl 1492R primers [72], complemented with 1µl dNTP, 1µl Dreamtap polymerase (Thermo Scientific), 5µl 10×Dreamtap buffer (Thermo Scientific) and 36µl H_2_O. The PCR reaction was processed on a thermocycler (Hybaid, Ashford, UK) with the cycling conditions in Table S10. PCR products were sequenced at Macrogen Europe (Amsterdam, the Netherlands). The 16S rRNA sequence were processed with MEGA 10.2.0 [73] and submitted to EzBioCloud 16S database [74] for taxonomy identification. We then mapped the 16S rRNA sequence of the isolates hyphosphere and bulk soil bacterial ASVs using VSEARCH [75] at 99% sequence similarity.

### Screening of mycorrhiza-associated bacteria for impact on plant growth

Prunella seeds were vapor-phase sterilized by exposure to chlorine gas for 4 h. The seeds were sown on agar-solidified half-strength Murashige and Skoog basal medium (Sigma Aldrich, St. Louis, MO, USA), with maximally 10 seeds per square Petri Dish (120×120mm, Greiner). Seeds were allowed to germinate and develop in a climate chamber under controlled conditions (short-day: 10h light/14h dark, 22°C). Two-week-old seedlings with roots of approximately ∼ 0.5 cm in length that were free of visible contaminations were used in our experiment.

River sand was autoclaved twice at 121℃ for 45mins and mixed thoroughly with OS in a ratio of 4:1 (w/w). *Devosia* sp. ZB163 (HB1), *Bosea* sp. ZB026(HB2), *Sphingopyxis* sp. ZB004 (HB3), *Achromobacter* sp. ZB019 (HB4), and *Microbacterium* ZB113 (HB5), *Arthobacter* sp. ZB074 (SB1), *Streptomyces* sp. ZB117 (SB2) and *Pseudomonas* sp. ZB042 (SB3) were streaked on ISP2 media and incubated at 28℃ for three days. A single bacterial colony was then suspended with a loop in 50 µl 10mM MgSO_4_, spread over a Petri-dish with ISP2 agar-solidified medium, and incubated at 28℃ overnight until the bacterial growth covered the full plate. Subsequently, 10ml of 10mM MgSO_4_ was added to the plates and the bacteria were suspended with a sterile spatula. The suspension was then collected in a 15-ml Greiner tube followed by a double round of centrifugation and resuspension of the pellet in 10 ml 10mM MgSO_4_. Finally, the suspensions of bacterial isolates were mixed through the sand/soil mixture to a final density of 3×10^7^ CFU/g of soil. Moreover, we inoculated a SynCom of 5 HB and a SynCom of 3 SB, both inoculated at a total density of 3×10^7^ CFU/g of soil. Soil for the control treatments received an equal amount of sterile 10mM MgSO_4_. For each treatment, we filled 11 replicate 60-ml pots, resulting in a total of 110 pots (10 treatments x 11 replicates). One *P. vulgaris* seedling was sown in each pot and plants were grown in a greenhouse for 9 weeks with 16h light/8h dark at 22°C. Each pot received 10 to 15ml of water three times a week. For the last three weeks, each plant was supplied with 15ml of ½ strength Hoagland (Table S5) solution once a week.

Shoots were cut at the soil surface, lyophilized and weighted. Plant roots were removed from the soil and rinsed in sterile water. A 1-cm-long fragment of rinsed root was cut, weighted and stored in 50% ethanol for mycorrhizal root colonization analysis. The colonization of mycorrhizae on plant roots was evaluated using the method outlined previously.

### Propagation of AM fungi for pot experiments studying the impact of hyphal associated bacteria on plant growth

We cultured Ri T-DNA-transformed carrot root organs on one side of a two-compartment petri dish at 26°C for 2 weeks and then inoculated the organs with spores of *Rhizophagus irregularis* MUCL43194 [76]. The root compartments were filled with modified Strullu and Romand (MSR; Duchefa Biochemie, NL) medium supplemented with 1% sucrose and the hyphal compartment were filled with MSR medium (Table S11). *R. irregularis* then was left to colonize the root organs for 3 months during which *R. irregularis* mycelium colonized the hyphal compartment of the Petri-dish and formed spores. *R. irregularis* spores were harvested by chopping the agar-solidified medium of the hyphal compartment into small pieces using a sterile scalpel and subsequently dissolving the medium in a sterile citrate buffer (Citric acid, 0.3456g/L; Sodium citrate, 2.4108g/L). Thousands of *R. irregularis* spores in citrate buffer were then transferred to sterile 1.5-ml Eppendorf tubes in 500-µl aliquots and stored at 4°C.

### Impact of *Devosia* sp. ZB163 and AM fungi on plant growth

Organic soil-sand mixture was autoclaved twice to remove the indigenous microbiota and was inoculated with *Devosia* sp. ZB163 in 10mM MgSO_4_ at a density of 3×10^7^ CFU/g of soil (*Devosia* treatment) or an equal volume of 10mM MgSO_4_ as mock control. Two-week-old Prunella seedlings were transplanted into 60-ml pots filled with both soil treatments. Half of the pots received 100 *R. irregularis* spores immediately prior to seedling transplantation (AM treatment). Eleven replicate pots were prepared for each of the 4 treatments (Control, *Devosia*, AM, and *Devosia* & AM) resulting in a total of 44 pots. Plants were allowed to grow under climate-controlled conditions at a light intensity of 200µE/m^2^/s with a 16h photoperiod for 8 weeks at 22°C. Each pot received 10 to 15ml of water three times a week. To determine the effect of N and P availability on plant growth, we conducted a complementary experiment with the same four treatments and 20 biological replicates, resulting in a total of 80 pots. Moreover, the plants were watered when appropriate, and for the experiment shown in Fig 7, plants were supplied with 5ml modified Hoagland solution without N or P (Table S5) once per week from week 6 onwards. Following the 8th week of cultivation, shoot weight, root weight, and mycorrhization were assessed as described above.

### N and P accumulation in plant leaves

Lyophilized Prunella leaves were first ground to powder. To determine P content, approximately 50 mg of powdered leaves were digested in 1ml HCl/HNO_3_ mixture (4:1, v/v) in a closed Teflon cylinder for 6h at 140℃. The P concentrations were determined colorimetrically using a Shimadzu UV-1601PC spectrophotometer [77]. The N concentrations were determined by dry combustion of a 3-4 mg sample with a Flash EA1112 elemental analyzer (Thermo Scientific, Rodano, Italy).

### Absolute quantification of *Devosia* sp. ZB163 on plant roots

To quantify the absolute abundance of *Devosia* sp. ZB163 on plant roots, we spiked root samples with 14ng DNA of *Salinibacter ruber*, an extremely halophilic bacterium that exists in hypersaline environments [40], but does not occur in our soil samples. Subsequently, the DNA of the root samples was extracted using the DNeasy PowerLyzer PowerSoil Kit (Qiagen, Hilden, Germany) following the manufacturer’s instructions. The 16S rRNA gene V3-V4 region was amplified following a two-step PCR using the primers 341F and 806R [69] and barcoding primers [78]. The amplified DNA was cleaned-up, quantified, normalized, pooled and subsequently sequenced on the Novaseq 6000 SP platform (2 × 250 bp) by Genome Quebec (Montreal, Canada). The raw sequencing data were demultiplexed, trimmed, dereplicated, and filtered for chimeras by DADA2 [79] in the QIIME2 environment (version 2019.07, https://qiime2.org/) [80]. Amplicon sequence variants (ASVs) were generated and annotated against the SILVA reference database (v132) [81]. ASVs assigned to mitochondria and chloroplast were removed. Since ASVs that are present in only a few samples may represent PCR or sequencing errors, we removed the ASVs that were present in ≤ 4 samples. Filtered ASV counts were constructed into an ASV table. The absolute abundance amount of detected *Devosia* sp. ZB163 DNA using the following formula.

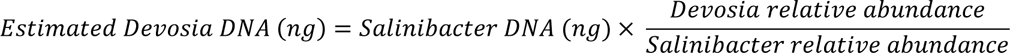

### *Devosia* genome sequencing

*Devosia* sp. ZB163 was cultured on ISP2 medium for 7 days at 28 ℃. DNA was extracted from a loop of bacterial cells using the MagAttract Microbial DNA Kit (Qiagen, Hilden, Germany) following the manufacturer’s instructions. The extracted DNA was amplified following the Hackflex protocol [82] followed by DNA purification using the AMPure XP clean-up (Beckman Coulter, High Wycombe, UK). The purified DNA was sequenced with Novaseq 6000 SP platform (2 × 250 bp) by Genome Quebec (Montreal, Canada). The raw sequencing data were trimmed with Cutadapt. Quality checked and assembly was performed using the A5-miseq pipeline [83].

### Genome analysis

*Devosia* sp. ZB163’s genome was annotated using prokka [84] and RAST [85]. Mining for orthologs of genes in the genomes of *Devosia* was performed using reciprocal BLASTp analysis. Genes were considered orthologs when the e-value was smaller than 10^−5^. Moreover, the whole *Devosia* genome was blasted against a *nifH* database [86] formatted for the dada2 pipeline [87].

### Bioinformatics

Sequence reads were processed in the Qiime2 environment (version 2019.07, https://qiime2.org/) [80]. We used the Demux plugin to assess paired-end sequence quality. The imported primer sequences were removed using Cutadapt [88]. The paired-end sequences were dereplicated and chimeras were filtered using the Dada2 denoise-paired script [79], which resulted in the identification of ASVs and a count table thereof. Fungal ITS2 sequences were further processed by filtering nonfungal sequences using ITSx [89]. 16S and ITS2 ASVs were taxonomically annotated employing a pre-trained naive Bayes classifier [90] against the SILVA (v132) [81] and UNITE (v8) [91] database, respectively. From this taxonomic annotation, 16S ASVs assigned as mitochondria and chloroplast were removed.

### Statistical analysis

All statistical analyses were conducted in R version 4.0.2 [92]. All bioinformatic files generated by Qiime2 were imported to R with Qiime2R [93]. Bray-Curtis distances were calculated by and visualized in principal coordinate analysis (PCoA) using the *Phyloseq* package [94]. Pairwise permutational analysis of variance (PERMANOVA) was performed using Adonis function in the Vegan package with 9999 permutations [95]. The visualization of microbial taxonomy and differentially abundant ASVs between sample types used ggplot2 [96] and Complex Heatmap package [97]. ASVs that are positively associated with hyphosphere, or soil microbiome were identified by R package *indicspecies* [37] and considered robustly enriched if their abundance was significantly higher in hyphal samples than both roots and soil samples as determined by one-way analysis of variance (ANOVA). The effect of microbial treatments on plant weight, AM fungi colonization rate, and plant nutrient uptake was assessed by one-way ANOVA and followed by the Tukey HSD test. Absolute abundance of *Devosia* sp. ZB163 was assessed for variation among treatments by ANOVA and followed by a Tukey HSD test. The correlation between *Devosia* sp. ZB163 absolute abundance and plant weight, AM fungi colonization, and plant nutrient uptake were assessed by simple linear regression.

## Author information

### Authors and Affiliations

Plant-Microbe Interactions, Department of Biology, Faculty of Science, Utrecht University, Padualaan 8, 3584 CH Utrecht, the Netherlands

Changfeng Zhang, Marcel G. A. van der Heijden, Bethany Kate Dodds, Thi Bich Nguyen, Jelle Spooren, Roeland L. Berendsen

Plant Soil Interactions, Division Agroecology and Environment, Agroscope, Reckenholzstrasse 191, CH-8046 Zürich, Switzerland

Marcel G. A. van der Heijden & Alain Held

Department of Plant and Microbial Biology, University of Zurich, Zollikerstrasse 107, CH-8008 Zurich, Switzerland

Marcel G. A. van der Heijden

Mycology, Earth and Life Institute, Université Catholique de Louvain, Louvain-la-Neuve, Belgium

Marco Cosme

Plants and Ecosystems, Biology Department. University of Antwerp, Belgium

Marco Cosme

### Contributions

M.G.A.v.d.H. initiated the research. C.Z., R.L.B., and M.G.A.v.d.H. conceived and designed the experiments. C.Z., B.K.D., T.B.N. collected the samples and performed the greenhouse experiments. C.Z., B.K.D., T.B.N., and J.S. isolated DNA from the collected samples and prepared the DNA libraries. C.Z. and A.H. isolated bacteria from fungal hyphae. C.Z. and T.B.N. identified the bacteria taxa. M.R.C. cultured the monoxenic mycorrhiza spores and provide suggestions for mycorrhiza inoculation. C.Z. analyzed the data. C.Z., M.G.A.v.d.H., M.C. and R.L.B. wrote the manuscript.

### Corresponding author

Correspondence to Roeland L. Berendsen

## Availability of data and material

The raw sequencing data of *Devosia* genome are deposited at the National Center for Biotechnology Information, GenBank database (https://www.ncbi.nlm.nih.gov/genbank/) by the accession PRJNA931835. The raw sequencing data of the amplicon reads are deposited at the European Nucleotide Archive (http://www.ebi.ac.uk/ena) by the study PRJEB59555.

## Acknowledgements

We thank Utrecht Sequencing Facility for providing sequencing service and data. We are grateful to Dr. Claire E. Stanley from Imperial College London, for providing suggestions on hyphal bacteria isolation. We thank Richard van Logtestijn and Rob Broekman from Vrije Universiteit Amsterdam for determining the N and P concentrations on Prunella leaves. We also thank Gijs Selten from Universiteit Utrecht for assembling the *Devosia* genome.

## Ethic declarations

### Ethics approval and consent to participate

Not applicable.

### Consent for publication

Not applicable.

