## Supplementary information for "A tripartite bacterial-fungal-plant symbiosis in the mycorrhiza-shaped microbiome drives plant growth and mycorrhization"

The following Supporting Information is available for this article:

Supplementary Figures

**Fig. S1. Effects of field management practices on soil microbial communities in Experiment I.**

**Fig. S2. Photo of root colonization in COMP3 of a representative mesocosm at the end of Experiment II.**

**Fig. S3. Effects of field management practices on soil microbial communities in Experiment I.**

**Fig. S4. Bacterial ASVs with significantly different abundance between hyphal and soil samples in Experiments I (A) and II (B).**

**Fig. S5. Isolation of AM-associated microbes using two strategies.**

**Fig. S6. Pearson's correlation between AM fungi root colonization (%) and plant P accumulation.**

**Fig. S7. Schematic representation of the wet sieving protocol used to sample hyphae from COMP 5 as described in the Methods section.**

**Fig. S8. Stereo microspore images of AM hyphae.**

Supplementary Tables

**Table S1. Effect of sample type on fungal and bacterial communities of experiment I.**

**Table S2. Effect of preceding soil management practices in the FAST experiment on microbial communities of root, hyphal and soils samples at the end of experiment I.**

**Table S3. Effect of the presence of plant on soil microbial communities.**

**Table S4. Effect of sample type on fungal and bacterial communities of experiment II.**

**Table S5. Hoagland solution ingredients.**

**Table S6. Overview of microbial genes involved in N metabolism for which orthologs were putatively found in the genome of *Devosia* sp. ZB163.**

**Table S7. Primers used for amplification of microbial ITS and 16S.**

**Table S8. Two step PCR cycling conditions for amplifying ITS, 16S.**

**Table S9. AM-associated bacteria isolation media.**

**Table S10. PCR cycling conditions for amplifying 16S.**

**Table S11. Modified Strullu and Romand (MSR) medium supplemented with 1% sucrose.**

**Additional file 1. Overview of the 144 bacteria isolated from hyphal samples.** This file contains Unique ID, taxonomy, FASTA sequence of the hyphal bacterial isolates.

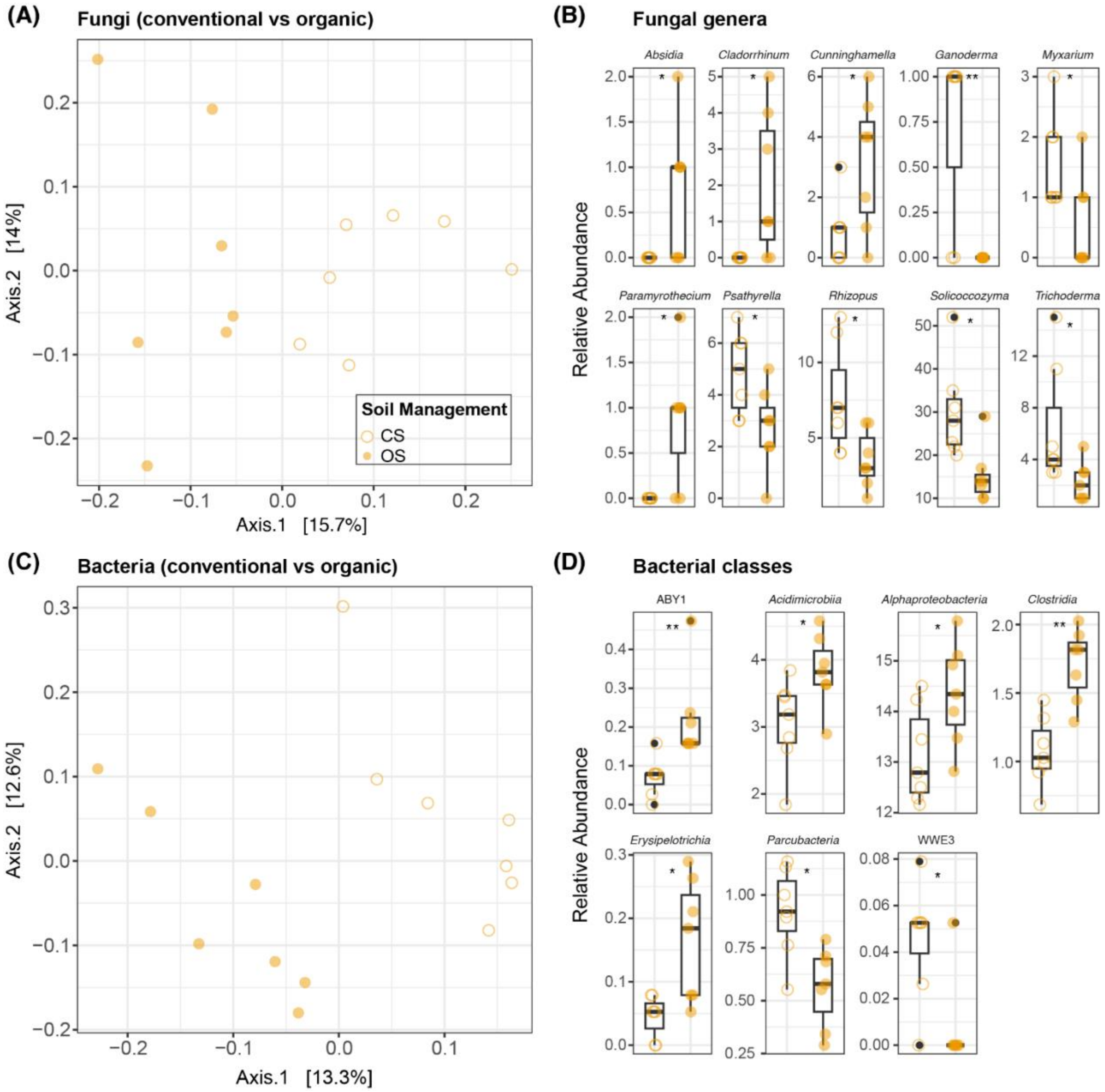


**Fig. S1. Effects of field management practices on soil microbial communities in Experiment I. (A)** PCoA of fungal communities using Bray-Curtis distances in CS and OS. **(B)** Relative abundance of fungal genera that are differentially abundant between CS and OS. **(C)** PCoA of bacterial communities using Bray-Curtis distances in CS and OS **(D)** Soil bacterial community differential abundant classes in CS and OS. Open orange circle stands for samples planted in CS, and closed orange circles stand for samples planted in OS. The number of y-axes in **(B)** and **(D)** show the percentage of relative abundance (%). The asterisk representing the *p*-value in **(B)** and **(D)** are determined by Wilcoxon text ($p^* < 0.05$, $p^{**} < 0.01$).

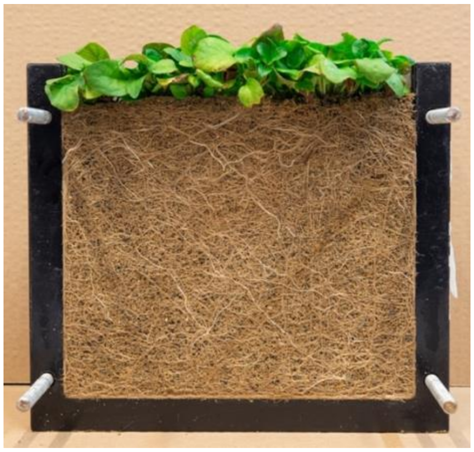

**Fig. S2. Photo of root colonization in COMP3 of a representative mesocosm at the end of Experiment II.**

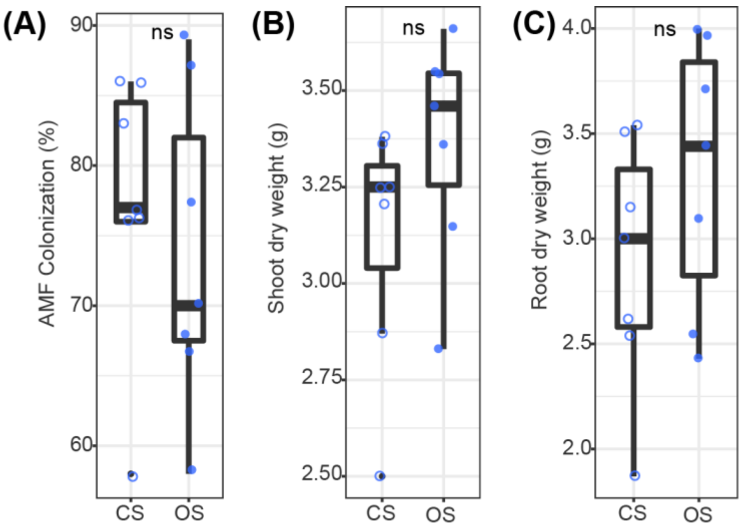


**Fig. S3. Effects of field management practices on soil microbial communities in Experiment I.** **(A)** PCoA of fungal communities using Bray-Curtis distances in CS and OS. **(B)** Relative abundance of fungal genera that are differentially abundant between CS and OS. **(C)** PCoA of bacterial communities using Bray-Curtis distances in CS and OS **(D)** Soil bacterial community differential abundant classes in CS and OS. Open orange circle stands for samples planted in CS, and closed orange circles stand for samples planted in OS. The number of y-axes in **(B)** and **(D)** show the percentage of relative abundance (%). The asterisk representing the p-value in **(B)** and **(D)** are determined by Wilcoxon text ($p^* < 0.05$, $p^{**} < 0.01$).

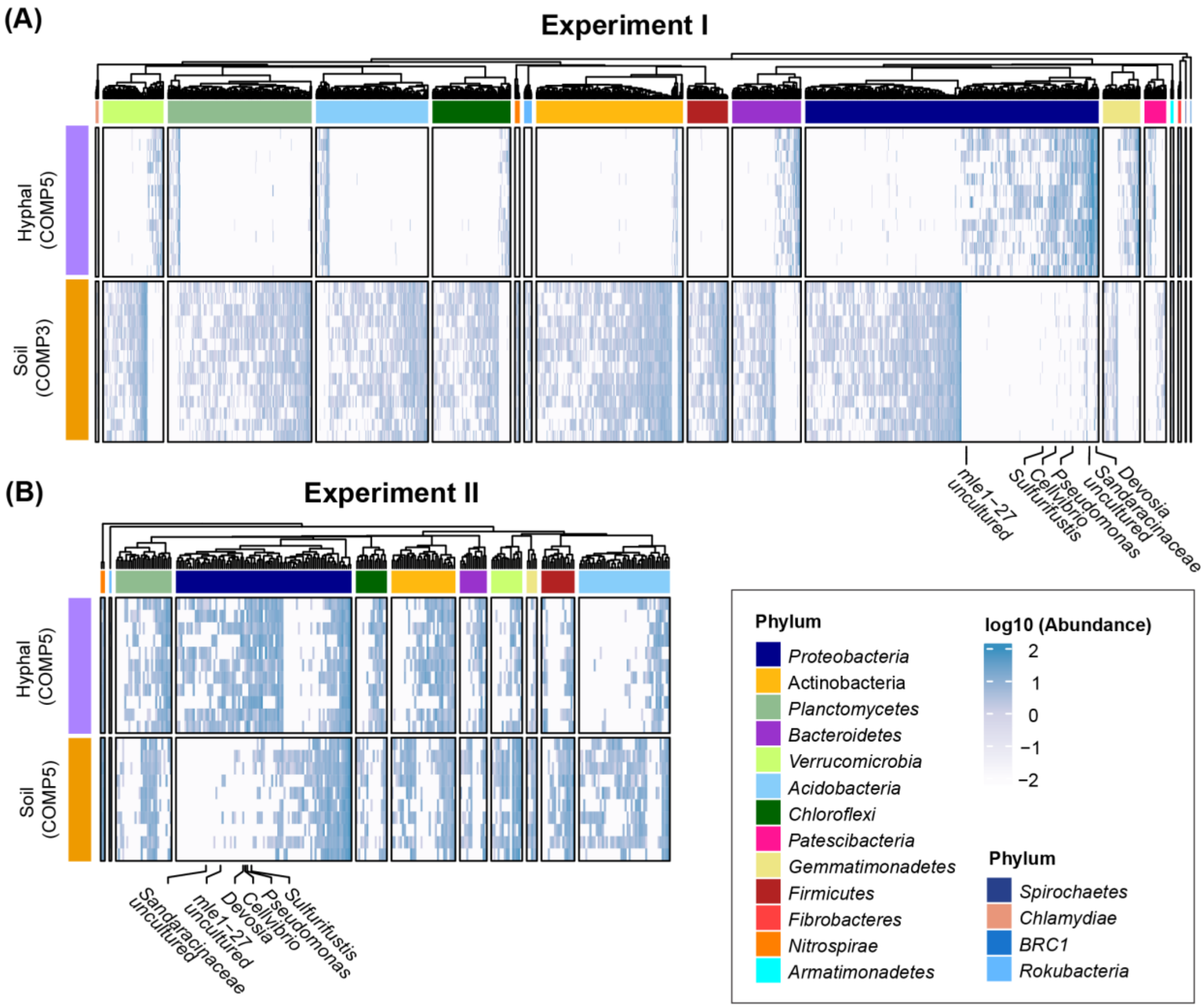


**Fig. S4. Bacterial ASVs with significantly different abundance between hyphal and soil samples in Experiments I (A) and II (B).** Heatmap shows log-transformed relative abundance of ASVs that significantly ($p < 0.05$) associate with either hyphal or soil samples. ASVs are ordered by phylogenetic distance and distinct phyla are indicated by vertical color bars on the left of the heatmap. Six consistently enriched bacterial ASVs are marked with their genus names or higher taxonomic rank when genus could not be identified. Bacterial phyla lower than 1% RA are not considered in the heatmap.

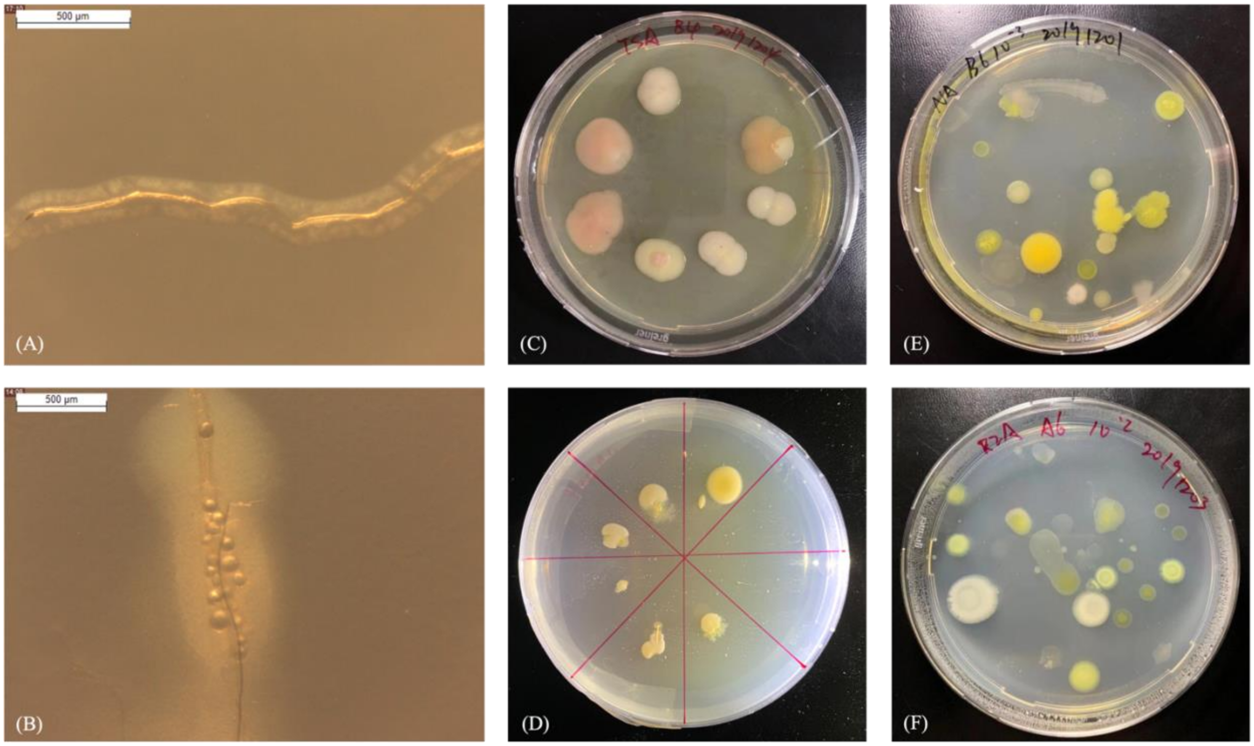


**Fig. S5. Isolation of AM-associated microbes using two strategies.** Stereo microscope images **(A)** and **(B)** show bacteria growing from mycorrhiza hyphae after 3 days of incubation. (Scale bar, 500 µm). **(C)** and **(D)** show bacterial colonies growing from mycorrhiza hyphae on agar plates after 20 days of incubation. **(E)** and **(F)** show bacterial colony-forming units grown from serial diluted hyphosphere samples after 20 days of incubation.

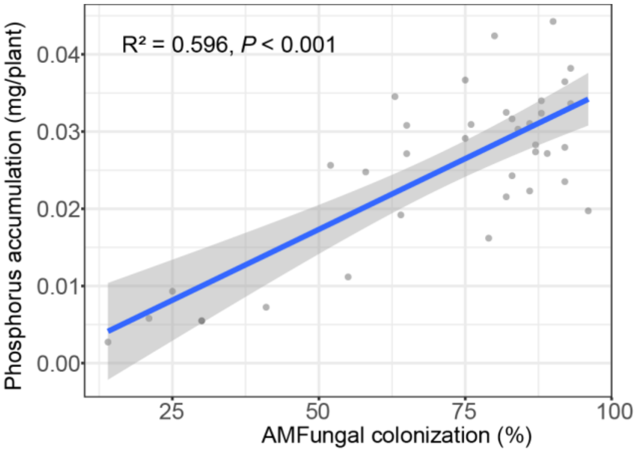


**Fig. S6. Pearson's correlation between AM fungi root colonization (%) and plant P accumulation.**

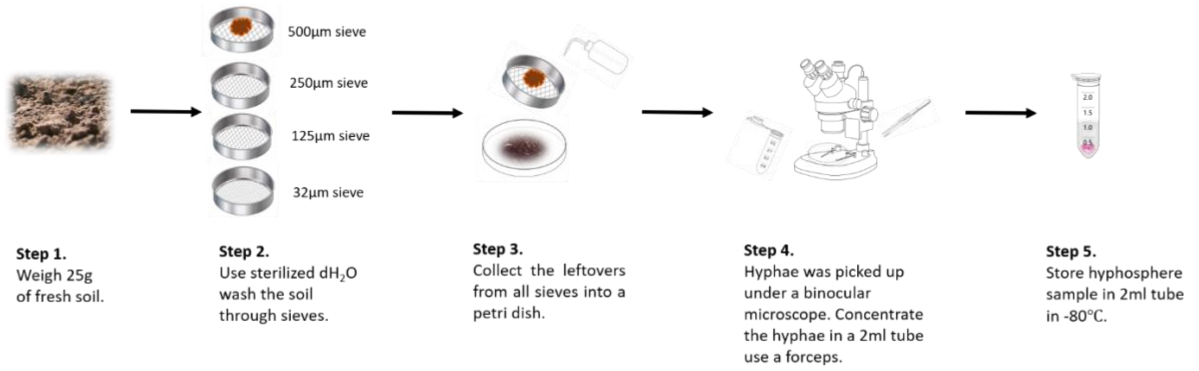


**Fig. S7. Schematic representation of the wet sieving protocol used to sample hyphae from COMP 5 as described in the Methods section.**

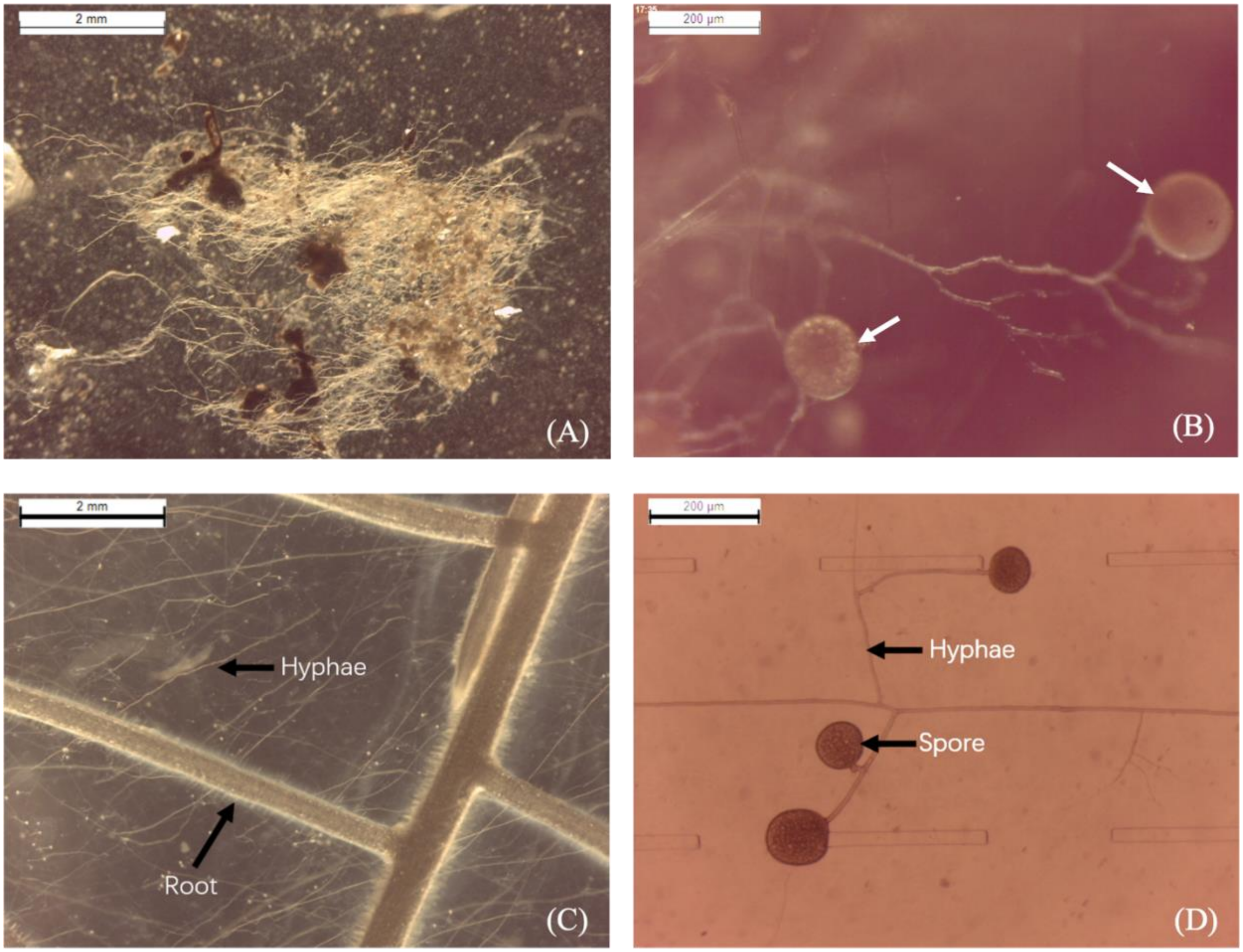


**Fig. S8. Stereo microspore images of AM hyphae.** Images of **(A)** hyphal sample acquired from COMP5 using the adapted wet sieving protocol (Scale bar, 2 mm). **(B)** AM spores within hyphal samples are indicated by arrows (Scale bar, 200 μm). **(C)** and **(D)** are references to AM hyphae cultured on an agar-solidified medium with chicory root organs. The scale bar of **(C)** and **(D)** are 2 mm and 200 μm respectively.

**Table S1. Effect of sample type on fungal and bacterial communities of experiment I.** Effect determined by pairwise PERMANOVA on Bray-Curtis distance with 9999 permutations.

| Sample type | Fungi | | | Bacteria | | |
|---|---|---|---|---|---|---|
| | F | $R^2$ | *p*-value | F | $R^2$ | *p*-value |
| **Root vs Hyphal** | **2.750** | **0.121** | **0.003*** ** | **16.06** | **0.391** | **<0.001*** ** |
| **Root vs Soil** | **22.563** | **0.485** | **<0.001*** ** | **16.018** | **0.381** | **<0.001*** ** |
| **Soil vs Hyphal** | **15.584** | **0.415** | **<0.001*** ** | **24.456** | **0.495** | **<0.001*** ** |

**Table S2. Effect of preceding soil management practices in the FAST experiment on microbial communities of root, hyphal and soils samples at the end of experiment I.** Pairwise PERMANOVA of the per samples type community is OS and CS on Bray-Curtis distance with 9999 **permutations.**

| CS vs OS | Fungi | | | Bacteria | | |
|---|---|---|---|---|---|---|
| | F | $R^2$ | *p*-value | F | $R^2$ | *p*-value |
| **Root** | 1.058 | 0.096 | 0.360 | 1.227 | 0.093 | 0.077. |
| **Hyphal** | 0.523 | 0.061 | 0.876 | 1.019 | 0.084 | 0.394 |
| **Soil** | **1.926** | **0.138** | **<0.001*** ** | **1.767** | **0.128** | **<0.001*** ** |

**Table S3. Effect of the presence of plant on soil microbial communities.** Fungal and bacterial communities are compared per compartment of experiment II by pairwise PERMANOVA on Bray-Curtis distance with 9999 permutations.

| Soil community in unplanted vs planted microcosm | Fungi | | | Bacteria | | |
|---|---|---|---|---|---|---|
| | F | $R^2$ | *p*-value | F | $R^2$ | *p*-value |
| **COMP1** | 0.093 | 0.074 | 0.217 | 1.155 | 0.081 | 0.156 |
| **COMP2** | **1.387** | **0.090** | **0.032*** | 0.948 | 0.068 | 0.637 |
| **COMP3** | **1.406** | **0.091** | **0.056.** | **1.537** | **0.099** | **0.003** ** |
| **COMP4** | 1.278 | 0.084 | 0.114 | **1.242** | **0.087** | **0.039*** |
| **COMP5** | 0.989 | 0.066 | 0.459 | 0.975 | 0.070 | 0.465 |

**Table S4. Effect of sample type on fungal and bacterial communities of experiment II.** Effect determined by pairwise PERMANOVA on Bray-Curtis distance with 9999 permutations.

| | Fungal | | | Bacteria | | |
|---|---|---|---|---|---|---|
| **Sample type** | F | $R^2$ | *p*-value | F | $R^2$ | *p*-value |
| **Root & Hyphal** | **7.182** | **0.274** | **<0.001*** ** | **11.389** | **0.375** | **<0.001*** ** |
| **Root & Soil** | **53.223** | **0.454** | **<0.001*** ** | **25.496** | **0.302** | **<0.001*** ** |
| **Soil & Hyphal** | **22.012** | **0.259** | **<0.001*** ** | **7.293** | **0.108** | **<0.001*** ** |

**Table S5. Hoagland solution ingredients.**

| Media | Compound | Amount |
|---|---|---|
| ½ Hoagland solution without N, P | **Macronutrients** | **Concentration (mM)** |
| | $K_2SO_4$ | 3 |
| | $CaSO_4$ ● $2H_2O$ | 2 |
| | $MgSO_4$ ●$7H_2O$ | 0.5 |
| | **Micronutrients** | **Concentration (µM)** |
| | KCl | 25 |
| | $H_3BO_3$ | 12.5 |
| | $MnSO_4$•$H_2O$ | 1 |
| | $ZnSO_4$•$7H_2O$ | 1 |
| | $CuSO_4$•$5H_2O$ | 0.25 |
| | $(NH_4)_6Mo_7O_{24}$•$4H_2O$ | 0.25 |
| | $C_{10}H_{12}FeN_2NaO_8$ | 10 |
| ½ Hoagland solution | **Macronutrients** | **Concentration (mM)** |
| | $KNO_3$ | 3 |
| | $(NH_4)H_2PO_4$ | 1 |
| | Ca(NO3)2•4H2O | 2 |
| | $MgSO_4$•$7H_2O$ | 0.5 |
| | **Micronutrients** | **Concentration (µM)** |
| | KCl | 25 |
| | $H_3BO_3$ | 12.5 |
| | $MnSO_4$•$H_2O$ | 1 |
| | $ZnSO_4$•$7H_2O$ | 1 |
| | $CuSO_4$•$5H_2O$ | 0.25 |
| | $(NH_4)_6Mo_7O_{24}$•$4H_2O$ | 0.25 |
| | $C_{10}H_{12}FeN_2NaO_8$ | 10 |

**Table S6. Overview of microbial genes involved in N metabolism for which orthologs were putatively found in the genome of *Devosia* sp. ZB163.**

| Gene | Location | Strand | Hits |
|---|---|---|---|
| ***nifU*** | scaffold_0_406316_406873 | + | NifU family protein [*Devosia* sp.] |
| ***fixK*** | scaffold_0_564243_563566 | - | CRP/FNR family N fixation transcriptional regulator |
| ***fixL*** | scaffold_1_81963_81343 | - | response regulator FixJ [*Devosia* sp. Root413D1] |
| ***fixJ*** | scaffold_1_83308_81950 | - | putative FixL oxygen regulated histidine kinase [uncultured bacterium 1062] |
| ***ureG*** | scaffold_3_209051_208419/ scaffold_3_594828_594178/ scaffold_0_761242_761880 | -/-/+ | urease accessory protein UreG [*Devosia rhizoryzae, Devosia oryziradicis*] |
| ***ureF*** | scaffold_3_210273_209473/ scaffold_3_597921_597244/ scaffold_0_760487_761230 | -/-/+ | urease accessory protein UreF [*Devosia rhizoryzae, Devosia oryziradicis*] |
| ***ureE*** | scaffold_3_210743_210270/ scaffold_3_598363_597914/ scaffold_0_759916_760527 | -/-/+ | urease accessory protein UreE [*Devosia* sp.] |
| ***ureD*** | scaffold_3_219092_218151/ scaffold_0_761880_762788 | -/+ | urease accessory protein UreD [*Devosia rhizoryzae, Devosia oryziradicis*] |
| ***Urease alpha subunit*** | scaffold_3_215267_213558/ scaffold_3_596522_594825/ scaffold_0_758160_759881 | -/-/+ | urease subunit alpha [*Devosia rhizoryzae, Devosia oryziradicis*] |
| ***Urease beta subunit*** | scaffold_3_217611_217306/ scaffold_3_597223_596528/ scaffold_0_757645_758115 | -/-/+ | urease subunit beta [*Devosia rhizoryzae, Devosia oryziradicis*] |
| ***Urease gamma subunit*** | scaffold_3_218142_217840/ scaffold_0_757305_757607 | -/+ | urease subunit gamma [*Devosia rhizoryzae, Devosia oryziradicis*] |
| ***UrtE*** | scaffold_3_219871_219095 | - | urea ABC transporter ATP-binding subunit UrtE [*Devosia rhizoryzae, Devosia oryziradicis*] |
| ***UrtD*** | scaffold_3_220730_219975 | - | urea ABC transporter ATP-binding protein UrtD [*Devosia rhizoryzae, Devosia oryziradicis*] |
| ***UrtC*** | scaffold_3_221902_220727 | - | urea ABC transporter permease subunit UrtC [*Devosia rhizoryzae, Devosia oryziradicis*] |
| ***UrtB*** | scaffold_3_223973_222219 | - | urea ABC transporter permease urea ABC transporter permease subunit UrtB [*Devosia rhizoryzae, Devosia oryziradicis*] |
| ***UrtA*** | scaffold_3_225350_224043 | - | branched-chain amino acid ABC transporter substrate-binding protein [*Devosia rhizoryzae, Devosia oryziradicis*] |

**Table S7. Primers used for amplification of microbial ITS and 16S.**

| Target gene | Primer pairs | Sequence |
|---|---|---|
| ITS | 5.8S Fun | AACTTTYRRCAAYGGATCWCT |
| | ITS4 Fun | AGCCTCCGCTTATTGATATGCTTAART |
| 16S | 341F | CCTACGGGNGGCWGCAG |
| | 805R | GACTACHVGGGTATCTAATCC |

**Table S8. Two step PCR cycling conditions for amplifying ITS, 16S.**

**First step**

| ITS | | | 16S | | |
|---|---|---|---|---|---|
| Temperature | Time | Cycles | Temperature | Time | Cycles |
| 96℃ | 2min | 1× | 95℃ | 3min | 1× |
| 94℃ | 30sec | | 95℃ | 30sec | |
| 58℃ | 40sec | 25× | 55℃ | 30sec | 25× |
| 72℃ | 2min | | 72℃ | 30sec | |
| 72℃ | 10min | 1× | 72℃ | 5min | 1× |
| 15℃ | Hold | - | 15℃ | Hold | - |

**Second step**

| ITS | | | 16S | | |
|---|---|---|---|---|---|
| Temperature | Time | Cycles | Temperature | Time | Cycles |
| 95℃ | 3min | 1× | 95℃ | 3min | 1× |
| 95℃ | 30sec | | 95℃ | 30sec | |
| 55℃ | 30sec | 10× | 55℃ | 30sec | 10× |
| 72℃ | 30sec | | 72℃ | 30sec | |
| 72℃ | 5min | 1× | 72℃ | 5min | 1× |
| 15℃ | Hold | - | 15℃ | Hold | - |

**Table S9. AM-associated bacteria isolation media.**

| Media | Compound | Amount/L |
|---|---|---|
| Trypic Soy Broth Medium | Casein | 17 g |
| | Soya peptone | 3 g |
| | NaCl | 5 g |
| | $K_2HPO_4$ | 2.5 g |
| | Dextrose | 2.5 g |
| | (Agar) | 20 g |
| | pH: 7.2 | |
| 1/10 TSA | Casein | 1.7 g |
| | Soya peptone | 0.3 g |
| | NaCl | 0.5 g |
| | $K_2HPO_4$ | 0.25 g |
| | Dextrose | 0.25 g |
| | (Agar) | 20 g |
| | pH: 7.2 | |
| Yeast Extract Manitol Medium | Yeast extract | 0.5 g |
| | Mannitol | 5 g |
| | $K_2HPO_4$ | 0.5 g |
| | $MgSO_4 \cdot 7H_2O$ | 0.2 g |
| | NaCl | 0.1 g |
| | (Agar) | 20 g |
| | pH: 7.0 | |
| Tap Water Yeast Extract Medium | Yest extract | 0.25 g |
| | $K_2HPO_4$ | 0.5 g |
| | (Agar) | 18 g |
| | Tap water to 1L | |
| | pH: 7.2 | |
| R2A medium | Casein acid hydrolysate | 0.5 g |
| | Yeast extract | 0.5 g |
| | Proteose peptone | 0.5 g |
| | Dextrose | 0.5 g |
| | Starch | 0.5 g |
| | Dipotassium phosphate | 0.3 g |
| | Magnesium sulfate | 0.024 g |
| | Sodium pyruvate | 0.3 g |
| | (Agar) | 15 g |
| | pH: 7.2 | |
| 1/5 R2A | Casein acid hydrolysate | 0.1 g |
| | Yeast extract | 0.1 g |
| | Proteose peptone | 0.1 g |
| | Dextrose | 0.1 g |
| | Starch | 0.1 g |
| | Dipotassium phosphate | 0.06 g |
| | Magnesium sulfate | 0.005 g |
| | Sodium pyruvate | 0.06 g |
| | (Agar) | 15 g |
| | pH: 7.2 | |
| Nutrient Agar | Peptone | 5g |
| | yeast extract | 3g |
| | NaCl | 5g |
| | Agar | 15g |
| | pH: 7.4 | |

**Table S10. PCR cycling conditions for amplifying 16S.**

| Step | Temperature | Time | Cycles |
|---|---|---|---|
| 1 | 94℃ | 5min | 1× |
| 2 | 94℃ | 1min | |
| 3 | 55℃ | 1min | 30× |
| 4 | 72℃ | 1min | |
| 5 | 72℃ | 10min | 1× |
| 6 | 12℃ | Hold | |

**Table S11. Modified Strullu and Romand (MSR) medium supplemented with 1% sucrose.**

| Component | Company | Amount/L |
|---|---|---|
| Strullu-Romand powder | Duchefa Biochemie (Haarlem, The Netherlands) | 0.594g |
| Sucrose | Sigma (St. Louis, Missouri, United States) | 10g |
| 0.152M $Ca(NO_3)_2$ | Merck (Darmstadt, Germany) | 10ml |
| Phytagel | Sigma (St. Louis, Missouri, United States) | 3g |
| $dH_2O$ | - | 976ml |

**Additional file 1. Overview of the 144 bacteria isolated from hyphal samples.** This file contains Unique ID, taxonomy, FASTA sequence of the hyphal bacterial isolates.
